# Rice Bean (*Vigna umbellata*) draft genome sequence: unravelling the late flowering and unpalatability related genomic resources for efficient domestication of this underutilized crop

**DOI:** 10.1101/816595

**Authors:** Tanushri Kaul, Murugesh Eswaran, Arulprakash Thangaraj, Arun Meyyazhagan, Mamta Nehra, Nitya Meenakshi Raman, Jyotsna Bharti, Gayacharan, Chandan Badapanda, Balamuralikrishnan Balamurali

## Abstract

Rice bean is a less well known and underutilized legume crop that has proved to be highly favourable due to its rich nutritional value in comparison with other members of the Vigna family. As an initiative to compose rice bean (*Vigna umbellata*) genomic resource, the size of 414 mega-base pairs with an estimate of 31276 highly confidential genes from 15521 scaffolds and functional coverage of 96.08% was sequenced from 30X coverage data from Illumina and PacBio platform. Rice bean genome assembly was found to be exquisitely close to V. angularis (experimental control/outgroup), *V. radiata* and *V. unguiculata,* however, *V. angularis* being the closest. Heuristically, the assembled genome was further aligned with 31 leguminous plants (13 complete genomes and 18 partial genomes), by collinearity block mapping. Further, we predicted similar discriminant results by complete CDS alignment. In contrast, 17 medically influential genomes from NIGMS-NIH, when compared with rice bean assembly for LCB clusters led to identification of more than 18000 genes from the entire selected medicinal genomes. Empirical construction of all genome comparisons revealed symplesiomorphic character in turn uncovering the lineage of genetic and functional features of rice beans. Signifiacantly, we found deserving late-flowering genes, palatably-indexed uncommon genes that regulate various metabolite pathways, related to abiotic and biotic stress pathways and those that are specific to photoperiod and disease resistance and so on. Further, we developed a repository for underutilised crop genome facility using D3.js at www.nicg.in. Therefore, the findings from this report addresses the genomic value of rice bean to be escalated via breeding by allied and applied approaches.

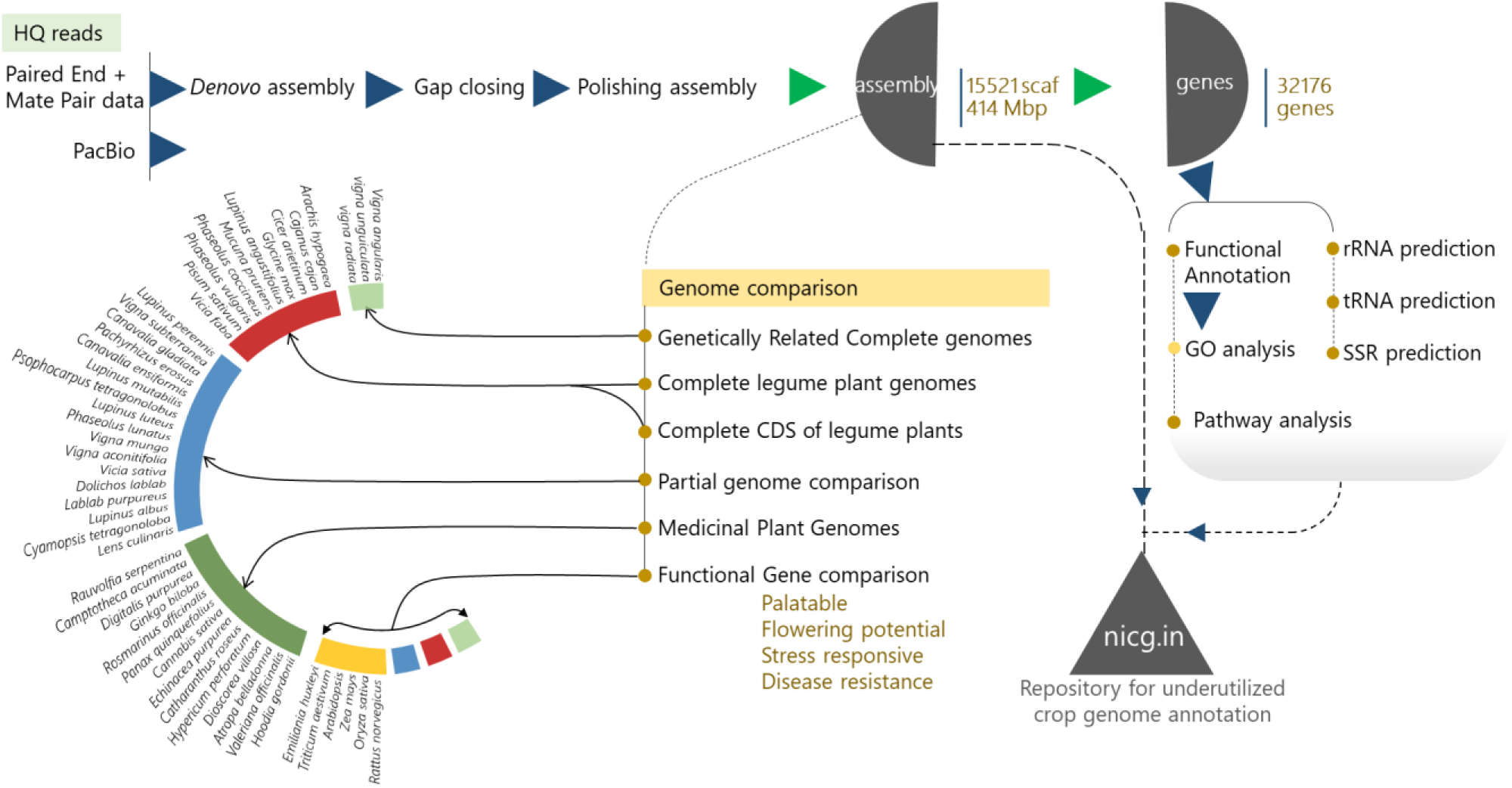

## Introduction

*Vigna umbellata* (Thumb.), is an orphan legume, multipurpose and lesser-known pulse with nutritionally rich crop. *Vigna* has been treated as an economically important genus that comprises of several domesticated species including *Vigna radiata* (mungbean), *Vigna angularis* (adzuki bean), *Vigna unguiculata* (cowpea) etc (Dahipahle *et al.,* 2019). Intriguingly, Anderson *et al.,* in 2009 report that the ricebean VRB3 that is an under-utilized crop has been nominated for multi-location testing amongst others and ranked no.1 with an average yield of 17.08 q/ha (Rana & Sood *et al.,* 2014). However, the dis-adoption of rice bean by farmers is linked to traits of landraces like late-flowering conditions [12h/d photoperiod (Takahashi *et al.,* 2015); 160-d-postanthesis (Joshi *et al.,* 2007)] and palatable gustations to humans (Basu and Scholten, 2013). The underlying molecular biology of these conditions is unexplored and elusive. CGIAR (Consultative Group for International Agricultural Research), IBPR (International Board for Plant Genetic Resources) and ICUC (International Centre for Underutilized Crops) organisations, form a network called FOSRIN (Food Security through Ricebean Research in India and Nepal), wherein Rice bean has been labeled as one of the future crops foreseen for domestication by farmers in marginal lands (Venkatesha 2012).

Despite the nutritional excellence of rice bean, the prevailing lack of awareness of its complete nutritional benefits means that rice bean can be categorized as an underutilized crop. Rice bean can become established and grow in various soil types. It is pest resistant and presents tremendous potential as a nutritious fodder and high-quality grain. The major drawbacks of this pulse crop of the kharif season include late flowering, indeterminate nature and tastelessness. Furthermore, there are meagre possibilities for the improvement of rice bean due to its predominant landraces, nominal modern plant breeding and limited seed supply. Consequently, the high diversity retained within its limited geographical distribution and the existence of few marketing channels mean that there is a great scope for the genetic improvement of rice bean. Despite the importance of rice bean as a multipurpose legume utilized for culinary purposes, the nutritious flour it produces, its use as a livestock feed and its efficiency as an accumulator of nitrogen in soil, the exploitation of this species in terms of productivity is meagre. There are several features of rice bean that require attention from geneticists before it can be widely adopted for standard consumption, including its high photoperiod sensitivity, late flowering period, high vegetative growth concerns, habit of twining (which makes harvest difficult), hard seededness and susceptibility to shatter (Smartt, 2012). Furthermore, the vulnerability of rice bean to pathogens results in heavy losses at harvest and reduced crop quality. The limited range of genomic tools for rice bean has impeded increases in its yield over a given period of time until now. There is an urgent need for crop improvement and the domestication of rice bean to generate new varieties with high-yield traits as well as pleasant organoleptic properties and reduced levels of antinutrients. Despite the fact that most of the available literature provides evidence of rice bean resistance to insect and aluminium toxicity (toxicity tolerance), no priority has thus far been given to understanding its flowering mechanism, palatability properties and disease resistance profiles, which are essential characteristics for achieving the maximum potential of this crop.

This study produced a detailed description of the nutritional attributes of this legume based on its genetic background. The results will increase awareness of the nutritional excellence of rice bean and improve genomics research on this crop. Our study involved the collection, screening, whole-genome sequencing, and genotype identification of rice bean and addressed the breeding of this crop with the aim of integrating high-protein and high-yield traits and determining rice bean growth in marginal areas and on hills and plains, starting from whole-genome sequencing with both the Illumina and PacBio platforms. A total of ∼69 Gb of data was generated from multiple sequencing platforms, including the Illumina (paired and mate pair) and PacBio platforms. Conventional genomic elements and novel genes for palatability domains, flowering potential and disease resistance were identified through the whole-genome sequencing of *Vigna umbellata* (Thumb.) Himshakti in different breeding lines, landraces and wild species to characterize genome-wide variations.

## METHODS

### Plant Material

The known phenotype background of *Vigna umbellata* (Thunb.) VRB3, seed was provided by ICAR-NBPGR (National Bureau of Plant Genetic Resources), New Delhi for WGS and PRR-2010-2, PRR-2011-2, RBHP-307, RBHP-104 and EC-14075 for transcriptome study of selected genes for flowering, palatability and stress response genes.

### DNA library preparation

A paired-end sequencing library was prepared using an Illumina TruSeq Nano DNA HT Library Preparation Kit (Kumar *et al.,* 2016). A 100-ng sample of g-DNA for the 350 bp insert size (2 libraries) and a 200 ng sample of g-DNA for the 550 bp insert size were fragmented with a Covaris instrument according to the manufacturer’s instructions. The covariance program for the 350 bp insert size and 550 bp insert size was calculated (Table 23).

**Table 23:**
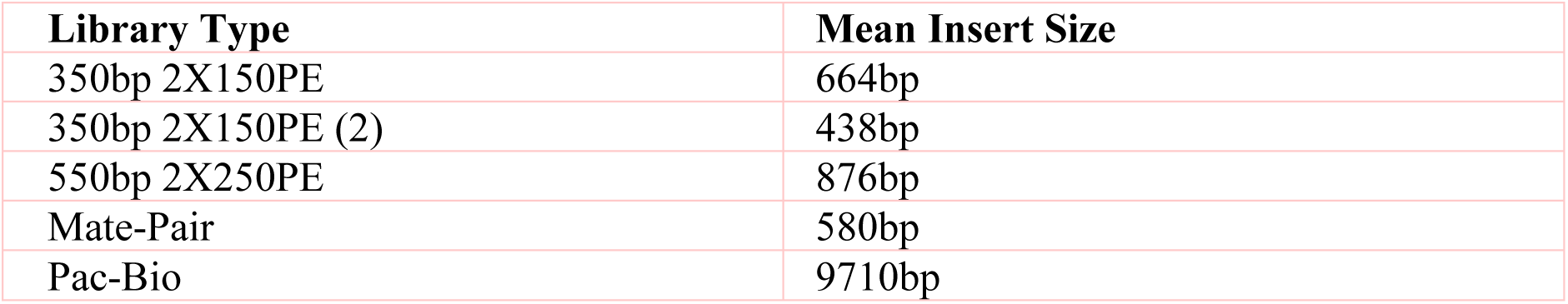
Complete library distributions concerning mean of insert size for each sequencing platform.

| Library Type | Mean Insert Size |
| --- | --- |
| 350bp 2X150PE | 664bp |
| 350bp 2X150PE (2) | 438bp |
| 550bp 2X250PE | 876bp |
| Mate-Pair | 580bp |
| Pac-Bio | 9710bp |

A mate-pair sequencing library was prepared using the Illumina Nextera Mate-Pair Sample Preparation Kit (Campbell *et al.,* 2016). One microgram of the gDNA was simultaneously fragmented and tagged with a biotinylated mate-pair junction adapter. An AMPure bead purification step was performed to clean up the PCR product and remove the smallest fragments (<300 bp) from the final library. Quantity and quality checks (QC) of the library were performed in a Bioanalyser 2100 instrument (Agilent Technologies) using a High-Sensitivity (HS) DNA chip.

### Comparative genome attributions

Genetic population structure analysis identified three distinct clusters in which the *Vigna umbellata* accession genome profiles and anchoring elements that correspond to functional features were mapped on the basis of the sequences of *Vigna angularis* (PRJNA328963) from the Beijing University of Agriculture, *Vigna radiata* (PRJNA243847) from Seoul National University and *Vigna unguiculata* (PRJNA381312) from the University of California, Riverside (Table 32), which are genetically and functionally orthologous to *Vigna umbelalta*. For all thirteen genomes selected for this study for the comparison of functional features, there are available complete genome profiles annotated via the Eukaryotic Genome Annotation Pipeline (EGAP). *Vigna angularis* was completely sequenced from 4 genome assemblies and 2 sequencing reads (Kang *et al.,* 2015), and the total genome size was approximately 455 MB, while the median total length of the *Vigna radiata* sequence (Kang *et al.,* 2014) was 459 mb, from 3 assemblies and 1 sequencing read, and that of *Vigna unguiculata* (Muñoz-Amatriaín *et al.,* 2017) was 607 mbp, from 2 genome assemblies.

**Table 32:**
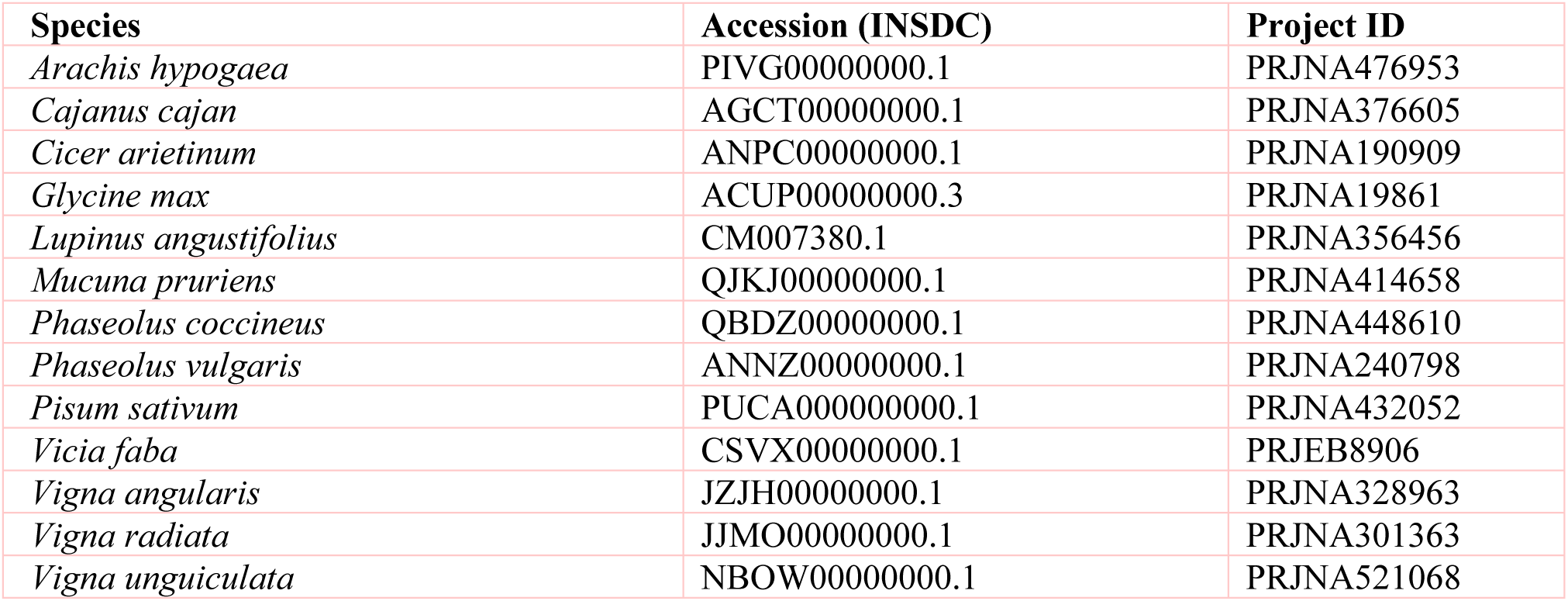
List of species Project ID and INSDC ID selected for genome comparison.

### Gene Predictions

Putative protein-coding genes were identified using the MAKER v2.31.9 pipeline (Campbell *et al.,* 2014), which identifies repeats, aligns transcripts and proteins to a genome, produces ab initio gene predictions and automatically synthesizes these data into gene annotations with evidence-based quality values. Three steps were followed to train the gene set. i) For evidence collection and ab initio gene prediction, in the first step, repeat regions were masked internally using RepeatMasker (Tarailo-Graovac and Chen, 2009), which screens DNA sequences for interspersed repeats and low-complexity DNA sequences. Then, the sequences were checked for their likely associations with genes by aligning the transcripts (using Exonerate) and proteins (using NCBI Blast) to the genome. In this case, we used the *Vigna radiata* protein and transcriptome dataset downloaded from NCBI (42284 proteins; GCF_000741045.1_*Vradiata*_ver6). The evidence was collected in GFF and fasta format files. All these steps were carried out in *(i)* the first round of MAKER execution to train the gene prediction software*. (ii)* In the second step, imperfect gene models from the first round of MAKER execution were used to the train gene predictor programs. Here, we used 2 gene prediction programs, SNAP [Semi-HMM-based Nucleic Acid Parser] (Korf I, 2004) and AUGUSTUS (Keller *et al.,* 2011). SNAP was trained using the MAKER bundled program, and AUGUSTUS was trained using BUSCOv3.0 (Waterhouse *et al.,* 2017). The outputs from both programs were used as training sets for the final MAKER run (second round of MAKER execution) to predict genes. *(iii)* For final gene prediction as a final step, polished genes are predicted using transcript and protein alignments and the HMM models generated in step (ii). MAKER takes all the evidence, generates “hints” about where splice sites and protein-coding regions are located, and then passes these “hints” to the gene prediction programs.

### Genome Library Preparation

Genomic DNA was isolated from *Vu* by using an Xcelgen CP Plant DNA isolation kit, and libraries were prepared using an Illumina TruSeq Nano DNA HT library preparation kit for insert sizes of 350 bp and 550 bp. The mean sizes of the libraries were 664 bp and 438 bp the 350 bp insert size and 876 bp for the 550 bp insert size. The library with an insert size of 350 bp was sequenced on the Illumina platform with 2 x 150 bp chemistry to generate ∼ 30 GB of data, while the library with an insert size of 550 bp was sequenced with 2 x 250 bp chemistry to generate ∼ 15 GB of data (Table 22). The mate-pair library was prepared from a ricebean sample using an Illumina Nextera Mate-pair Sample Preparation Kit. The mean size of the library was 580 bp. The library was sequenced (2 x 150 bp chemistry) to generate ∼ 10 GB of mate-pair data. The Pac-Bio library was prepared and sequenced using the PacBio sequencing platform to generate 6.5 GB of data.

**Table 22:**
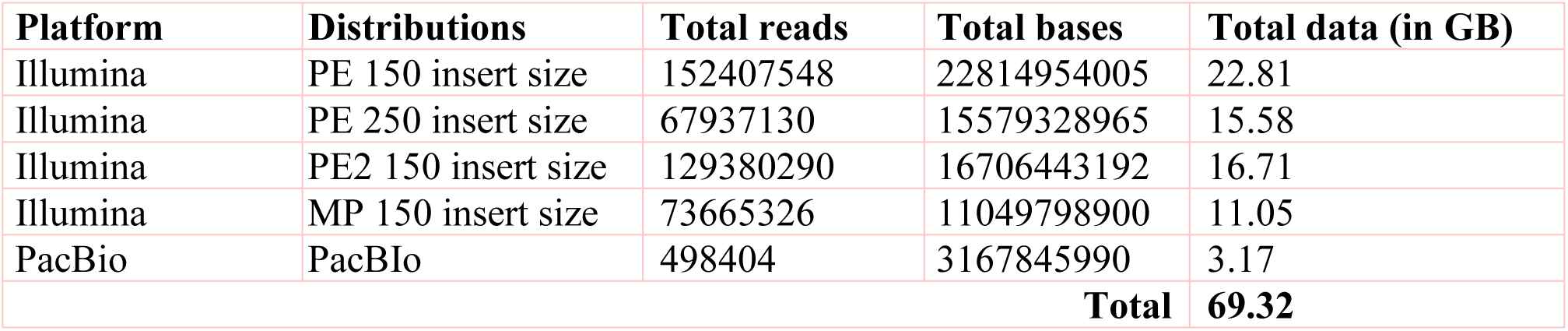
Complete library distributions of Vigna umbellata whole genome sequencing platform, total reads and total basepairs.

| Platform | Distributions | Total reads | Total bases | Total data (in GB) |
| --- | --- | --- | --- | --- |
| Illumina | PE 150 insert size | 152407548 | 22814954005 | 22.81 |
| Illumina | PE 250 insert size | 67937130 | 15579328965 | 15.58 |
| Illumina | PE2 150 insert size | 129380290 | 16706443192 | 16.71 |
| Illumina | MP 150 insert size | 73665326 | 11049798900 | 11.05 |
| PacBio | PacBio | 498404 | 3167845990 | 3.17 |
| **Total** |  |  |  | **69.32** |

### Alignment and tree building

Geneious Prime (v2019.1) used for the complete pre and post-assembly annotation process. Paired reads were trimmed further using BBDuk (Kim and Daadi, 2019) in Geneious to remove low-quality bases (Q < 20) and discard reads shorter than 10 bp. The assembled reads were then used for de novo assembly to generate maps using the Geneious *denovo* assembler. The consensus sequences of the scaffolds responsible for flowering, palatability, disease resistance and stress response genes were extracted from all selected species and aligned using the MAFFT (De Vega *et al.,* 2015) plugin. Third-party plugins such as Mauve aligner (Darling, Mau and Perna, 2010), tfscan, TadPole, RepeatMasker, and the customized local search library, extended reassembly aligner, and maximum likelihood Geneious plugins were used. Domain trees were estimated using the Maximum Likelihood (ML) plugin.

The tRNAscan-SE tool v1.3.1 (Varshney *et al.,* 2011) was used for the identification of selenocysteine tRNA derived genes, repetitive elements and pseudogenes from the assembled genome. A standalone version of tRNAscan-SE was used for the detection of tRNAs. Sequences encoding tRNAs were identified by tRNAScan-SE, which resulted in 577 tRNAs. Sequences encoding rRNAs were identified by RNAmmer (Zhuang *et al.,* 2019), which resulted in 11 rRNAs.

Using the MISA program, highly polymorphic and ubiquitous SSR markers were identified from the assembled genome. The assembled genome from both samples was searched for SSRs using the MISA tool (Singh *et al.,* 2011). A total of 67,523 SSRs were identified. Among the 67523 SSRs identified, 55699 exhibited a 150 bp flanking region that could be validated through PCR.

The BBH (bi-directional best hit) option was used to assign KO terms. Pathway analysis was carried out for assembled sequence using KAAS server (KEGG Automatic Annotation Server) By screening Bi-Directional best Hit. The genes were enriched in different functional pathway categories, predominantly related to Metabolism, Genetic Information Processing, Environmental Information Processing and Cellular Processes (Mochida *et al.,* 2016). A total of 676 genes contributed to the activity of the Carbohydrate Metabolism pathway, with the next-largest group of 657 genes was being related to Signal Transduction pathway.

GO annotations were obtained for the annotated genes using Blast2GO Command Line v-1.4.1. BLAST2GO PRO (Jain *et al.,* 2013) was used to annotate the functional genomic set for the assembled dataset. Gene Ontology terms fall under three categories: Biological Processes, Molecular Function and Cellular Components. Some of the genes were assigned by more than one category. A total of 11,204 terms fell under Biological Process, 13,097 under Molecular Function and 9,053 under Cellular Component.

To plot syntenic locally collinear blocks from Mauve alignment, the recently developed advanced real-time visualization tool Circa was used. Both the uni-dimensionality and connectivity of the two genomic datasets were plotted with Circa (OMGenomics, 2019).

Mauve Aligner: Multiple copies of the gene library were generated and aligned for comparison with all three reference genomes (Cheon *et al.,* 2019). The consistency or variation among the reference genomes was determined to compute locally colinear blocks.

PlasMapper generates plasmid maps along with annotations. It provides multiple predictions, such as replication origin, promotor, terminator, selectable marker, regulatory sequence affinity tag, miscellaneous feature, report gene and restriction site predictions (Shi *et al.,* 2017). The *Vigna umbellata* draft assembly was subjected to identify these elements to report additional genome-editing scaled tools.

d3js | DataDriven Documents: D3.js is a JavaScript library for manipulating documents based on genetic data. D3 helped us to obtain a dynamic view of the data using HTML, SVG, and CSS. D3’s emphasis on web standards provides the maximum capabilities of modern browsers without being tied to a proprietary framework, combining powerful visualization components and a data-driven approach for DOM manipulation (Garg *et al.,* 2017). The complete draft genome profile has been plotted using Gbrowse at www.nicg.in and is reported as “The Under-utilized Crop Genomic Facility” by NIC, ICGEB. New Delhi.

## RESULTS

### De novo assembly and Genome Statistics

With a benchmark of 69.32 GB of total raw data, 414 Mbps of reads were assembled using the Allpath LG (APLG) assembly pipeline (Kang *et al.,* 2015). To address the high-confidence draft of ricebean, the assembled reads from APLG were subjected to gap closure using PGJelly (Peng *et al.,* 2019). A total of 17,601 scaffolds were produced prior to gap closure and post-polishing with 3 GB of PacBio data. We obtained 15,521 final scaffolds and 36,461 contigs. A total of 41447 and 39,387 contigs were reported during the process of gap closure. The length of the total assembled reads was 387,954,823 bp; post-improvisation, a total of 414,706,990 bp was obtained from PBjelly, which aligns long sequencing reads. The average scaffold size was 26,719 from the final assembled reads (Table 1). The N50, N75 and N90 metrics were applied to weigh the quality of the genome assembly (Fig 1). The scaffold size distributed as 50% of the total was 78,112 bp; 75% of the total was 33,596 bp; and 90% of the total was 14,991 bp. The total no. of scaffolds for all three summation statistics were 7,652, 12,330 and 12,767 for N50, N75 and N90, respectively. The size of the N50 distribution is 207,323,106 bp (Table 2). The maximum scaffold size of the assembled sequences was 1,043,617 bp. The minimum scaffold size was 886 bp.

**Fig 1:**
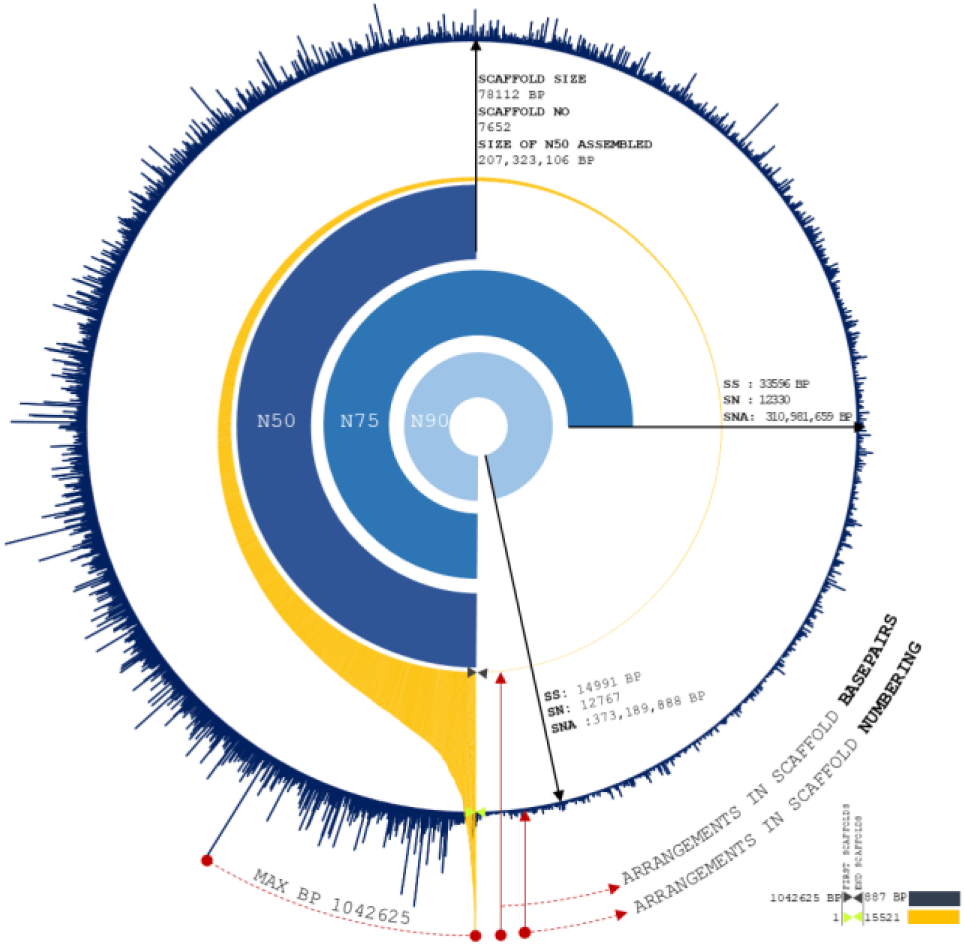
Assembly summation statistics : First layer (Blue) exhibit the distribution of scaffold number orderly over completeness of 15221 scaffold reads. Second layer (Orange) are the arrangement of scaffolds with respect to basepair length. Internal three layers are representing N50, N75 and N90 statistic of our assembly with scaffold size, scaffold position and size of summation statistics.

**Table 1:** Allpath-LG gapclosure and Pbjelly gapclosure comparison of Summation and assembly statistical composition from rice bean genome assembly.

| Statistics composition | Allpath-LG (primary) | Allpath-LG-Gapclosure | PBjelly | Pbjelly-gapclosure |
| --- | --- | --- | --- | --- |
| No. of scaffolds | 17601 | 17601 | 15521 | 15521 |
| No. of contigs | 41447 | 39387 | 36461 | 36241 |
| Total genome length including gaps | 389810950 (bp) | 387954823 (bp) | 414706990 (bp) | 414646213 (bp) |
| Total genome length without gaps | 301496373 (bp) | 351493463 (bp) | 382473710 (bp) | 387569263 (bp) |
| Average scaffold size including gaps | 22147 | 22042 | 26719 | 26715 |
| Scaffold N50 | 65704 | 65252 | 78103 | 78112 |
| Maximum scaffold size | 563916 (bp) | 564527 (bp) | 1043617 (bp) | 1042625 (bp) |
| Minimum scaffold size | 882 (bp) | 882 (bp) | 886 (bp) | 886 (bp) |

**Table 2:** Scaffold summation(N50, N75 and N90) distribution of rice bean assembled data.

|  | Scaffold Size | Scaffold No | Size of Assembly |
| --- | --- | --- | --- |
| N50 | 78112 | 7652 | 207323106 |
| N75 | 33596 | 12330 | 310981659 |
| N90 | 14991 | 12767 | 373189888 |

### Length distribution

*Assembly:* A total of 779 scaffolds were between >8000 and ≤10000. A total of 625 scaffolds >6000 and ≤8000, 803 scaffolds >4000 and ≤6000, 1519 scaffolds >2000 and ≤4000, 4095 scaffolds >1000 and ≤2000, 256 scaffolds >900 and ≤1000 and 10 scaffolds >300 and ≤900, 256 from >900 and ≤1000 (Table 3). Significantly, more than 7343 assembled scaffolds were greater than 10000 bp in length, which increases the possibility of understanding gene conservation and diversity in comparison with closely and distantly related species (VanBuren *et al.,* 2015) (Fig 2). *Genes predicted:* The maximum length of the genes involved in the prediction analysis was 15628 bp. A total of 31276 genes were predicted from 15521 scaffolds. Most of the genes predicted ranged between > 1000 and ≤ 2000 or >300 and ≤ 1000 (11442 and 111476 genes, respectively). A total of 8344 genes were distributed with the higher ranges of >8000 and ≤10000, >6000 and ≤8000, >4000 and ≤6000, and >2000 and ≤4000 (Table 4). Fourteen genes were predicted to present a size >10000.

**Fig 2:**
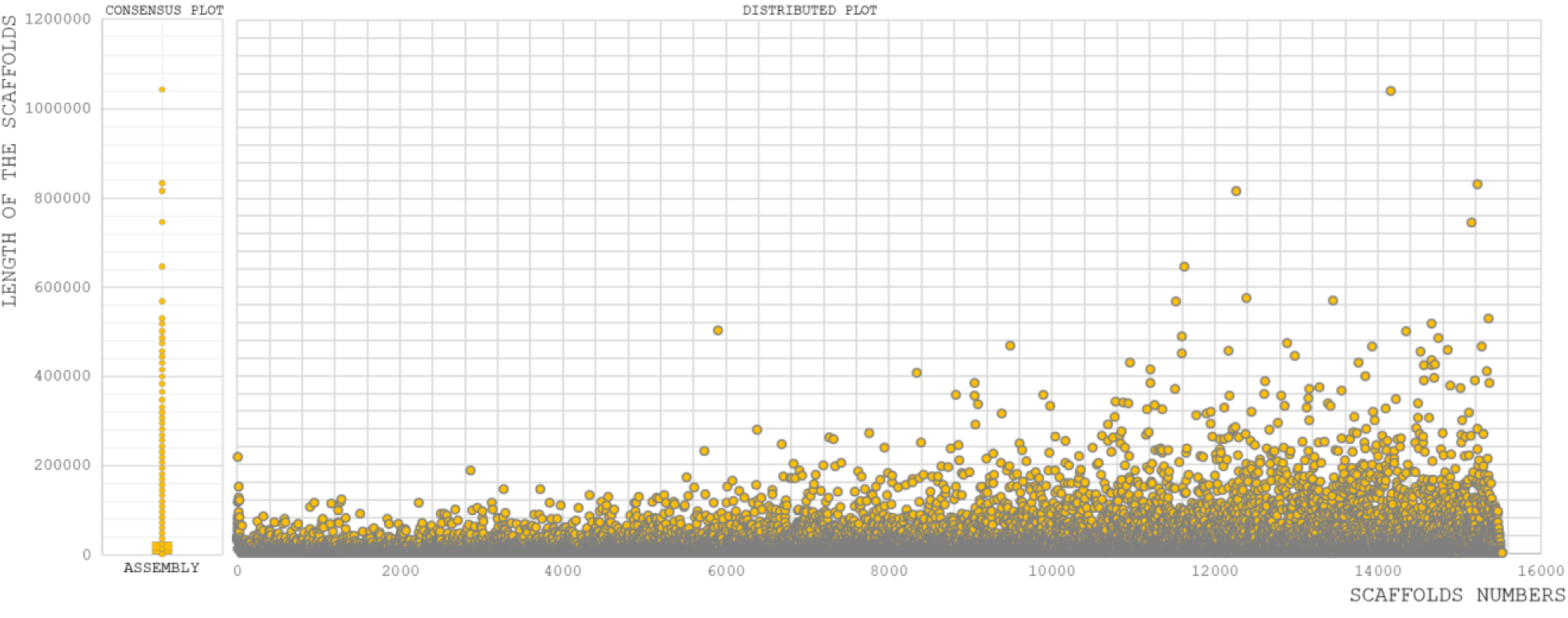
Sequence assembly length distribution coordinated from scaffold numbers to length of the sequence in each scaffold.

### Comparative Genome Statistics

Empirical distributions were studied by aligning the assembled reads with *Vigna angularis*(adzukibean)*, Vigna radiata*(mungbean), and *Vigna unguiculate* (cowpea). During the functional annotation of genes from the assembled scaffolds, 80% of genes were annotated to *Vigna angularis* (Tomooka, Norihiko. *et al.,* 2009), which is closely related to *Vigna radiata* and *Vigna unguiculate* (Fig 3). For *Vigna angularis*, 443 mbps was assembled from 290 GB of raw data, whereas for ricebean, 15521 scaffolds were assembled from 414 mbps (Kang *et al.,* 2015), and there is a high-confidence draft report from 30X-coverage library sequence (Table 5). The total numbers of scaffolds from mungbean and adzukibean were 2,748 and 3,883, respectively, and for ricebean 15,521 scaffolds predicted 31,276 genes.

**Fig 3:**
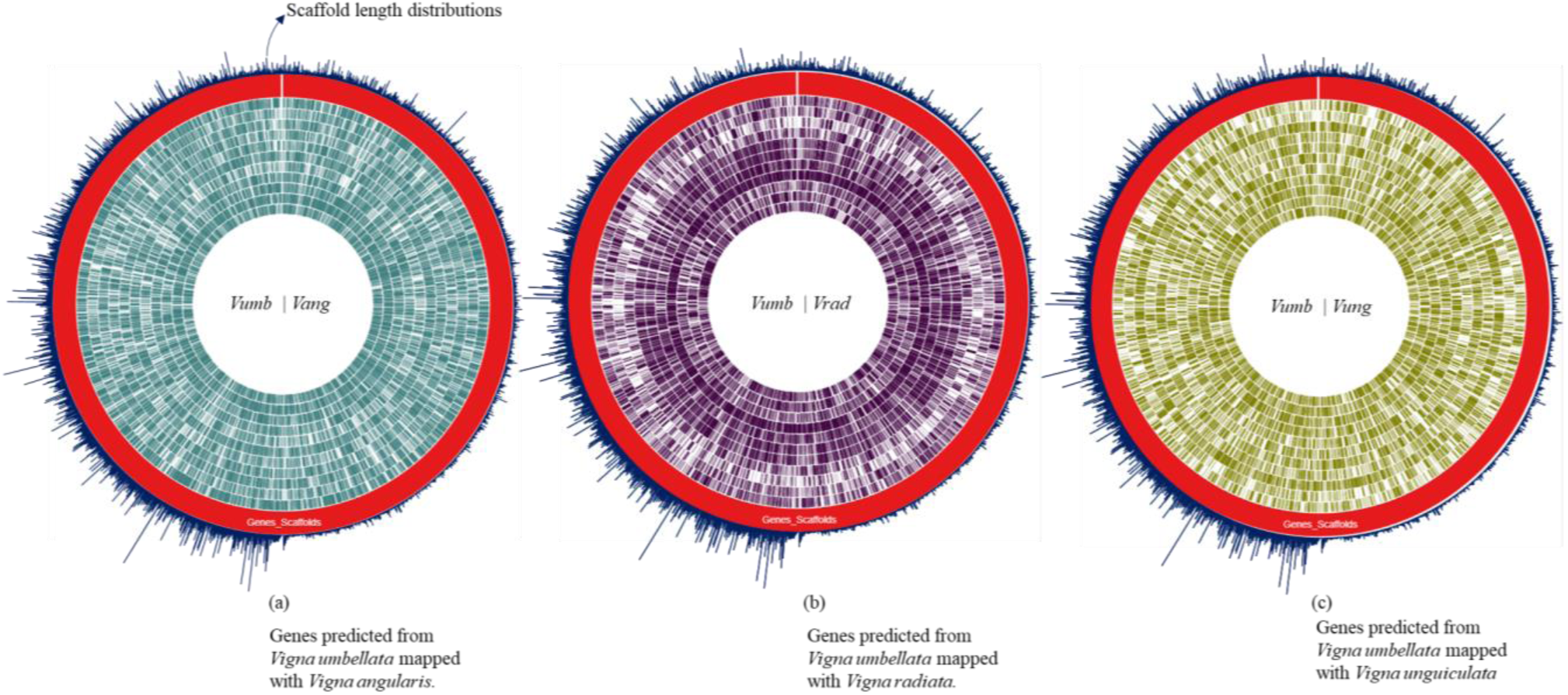
Complete predicted genes of Vigna umbellata mapped for Chromosomal specific comparison for the proximity of gene scaffold mapping to its reference. (a) Rice bean mapped syntenic region average of 82 % spanned for its gene specificity with adzuki bean. (b) Rice bean with mung bean having relation of each chromosomes of about 72 %. (c) Rice with cowpea mapping found 62 % of functional clusters with micro break from its evolution.

**Table 5:** Summation comparison of rice bean, mung bean and adzuki bean concerning total sequenced data and N50 scaffold distribution

| Statistics composition | *Vigna umbellata* | *Vigna radiata* | *Vigna angularis* |
| --- | --- | --- | --- |
| Total sequencing data | 69 Gb | 320 | 291 Gb |
| No. of scaffolds | 15521 | 2748 | 3883 |
| No. of contigs | 36241 | 25922 | 36516 |
| Tot. genome len. including gaps | 414646213 (bp) | 463143055 (bp) | 443436934 (bp) |
| Scaffold N50 | 78112 | 1516 | 703 |
| Minimum scaffold size(in bps) | 886 | 1000 | 1000 |

### Alignment read distribution for Comparative Genome annotations

The total length of the predicted genes was 49,811,559 bp (49mbp) from 414,646,213 bp (414mbp) of assembled reads. The predicted genes were mapped with three selected reference genome sequences: 105,415,055 bp from adzukibean, 92,715,800 bp from mungbean and 81,760,762 bp from cowpea were mapped to explore genes of ricebean (Table 6).

**Table 6:** Base pair distribution of Assembled reads: 31276 genes predicted from the assembly, bp of sequence mapped between genes predicted from rice bean and complete genome of adzuki bean, between genes predicted from rice bean and mung bean and between genes predicted from rice bean and cowpea.

| Read Profile | Length |
| --- | --- |
| Assembly reads | 414646213 (bp) |
| Genes predicted | 49811559 (bp) |
| Genes Predicted vs Vang bp | 105415055 (bp) |
| Genes Predicted vs Vrad bp | 92715800 (bp) |
| Genes Predicted vs Vung | 81760762 (bp) |

A total of 41,525 reads; predicted gene fragments were aligned with adzukibean, corresponding to 21,006 genes. A total of 32,657 and 16,614 reads were aligned with mungbean and cowpea, respectively, corresponding to 21657 and 20664 genes (Table 7).

**Table 7:** Total read count distribution of adzuki bean, mung bean and cowpea mapped with assembled reads. Total read count distribution of genes mapped from assembly with adzuki bean, mung bean and cowpea

| Read Alignment Profile | Counts |
| --- | --- |
| Total read aligned with Ref Vang | 41525 |
| Total read aligned with Ref Vrad | 32657 |
| Total read aligned with Ref Vung | 16614 |
| Total read aligned for gene annotation with Ref Vang | 21006 |
| Total read aligned for gene annotation with Ref Vrad | 21657 |
| Total read aligned for gene annotation with Ref Vung | 20664 |

### Gene Profile and Protein-coding regions

Out of the 31,276-total mapped ricebean genes, orthologous loci were found for 16892 genes (Fig 4) by gene mapping to adzukibean, 19,640 loci for mungbean and 17989 loci for cowpea (Table 8). Taking the positions of the CDS as functional regions for understanding the total composition of protein-coding genes, we found regions corresponding to non-coding RNA, which makes the calculation of *Vumb* genome accessibility for further domestication purposes easier. A total of 2 mbps of mapped genes from *Vang,* 5 mbps *from Vrad* and 9 mpbs from *Vung* were screened as non-coding RNA content.

**Fig 4:**
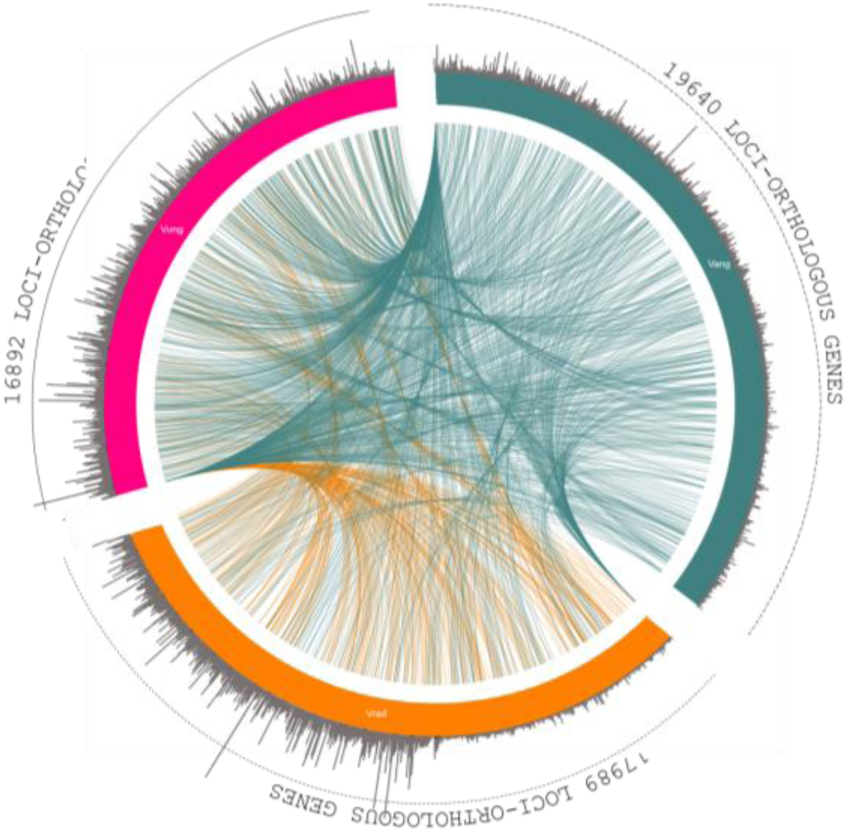
Orthologous synteny gene map comparison between Vang, Vrad, Vung with genes mapped from assembly. Green colour is adzuki bean mapping with 19640 loci, Orange colored mung bean mapping; 17989 loci. 16892 loci for cow pea-pink colored.

**Table 8:** Total read counts of total genes predicted from the assembly, read count of orthologous genes compared between mapped genes from assembly and complete reference genome of Vigna angularis; Read count of orthologous genes aligned between genes mapped with adzuki bean versus complete reference genome of adzuki bean, between genes mapped with mung bean versus complete reference genome of mung bean, and between genes mapped with cowpea versus complete reference genome of cowpea; CDS read counts between genes predicted and complete reference genome of adzuki bean, mung and cowpea.

| Gene Profile | Count |
| --- | --- |
| Total Genes Predicted | 31276 |
| Orthologous genes - Genes Predicted Vs Vang | 26001 |
| Orthologous genes - ( Genes Predicted vs Vang ) Vs Ref_Vang | 16892 |
| Orthologous genes - ( Genes Predicted vs Vrad ) Vs Ref_Vrad | 19640 |
| Orthologous genes - ( Genes Predicted vs Vung ) Vs Ref_Vung | 17989 |
| CDS match Genes Predicted with Ref_Vang | 8308 |
| CDS match Genes Predicted with Ref_Vrad | 8985 |
| CDS match Genes Predicted with Ref_Vung | 8515 |

#### tRNA operons

Codon usage biases and the use of identical anti-codons for tRNA pseudogenes that are caused by retrotransposon-driven repetitive elements (Gibbs RA, *et al.,* 2004) were assessed by using tRNAscan-SE with predetermined parameters (Int Cutoff = −30) for the assembled reads. A tRNA scan screened 577 tRNA operons that were positioned in intronic spacers and bound regions. The lowest covariance model score (Inf Score) calculated was 20.3, and the highest was 92.3. Searches were performed from beginning region to the end region of tRNAs and intronic spacers. The distribution of the amino acid plots with respect to codon occurrence was calculated. Arg-Leu-Ser showed the maximum frequency among the predicted codons. According to the standard genetic code representing direct amino acid synthesis, out of 6 codons, 5 codons were detected from scaffold inputs for arginine, leucine, and serine. Out of 4 codons, 3 codons were detected for Ala-Gly-Pro-Thr-Val; 2 codons were detected for Ile; 2 codons were detected for Asp-Gln-Glu-Lys; 2 codons were detected for Asn, Cys, His, Phe & Tyr; and 1 codon was detected for Met and Trp (Table 9). The total tRNA genes are summarized in the supplementary table (Fig 5).

**Table 9:**
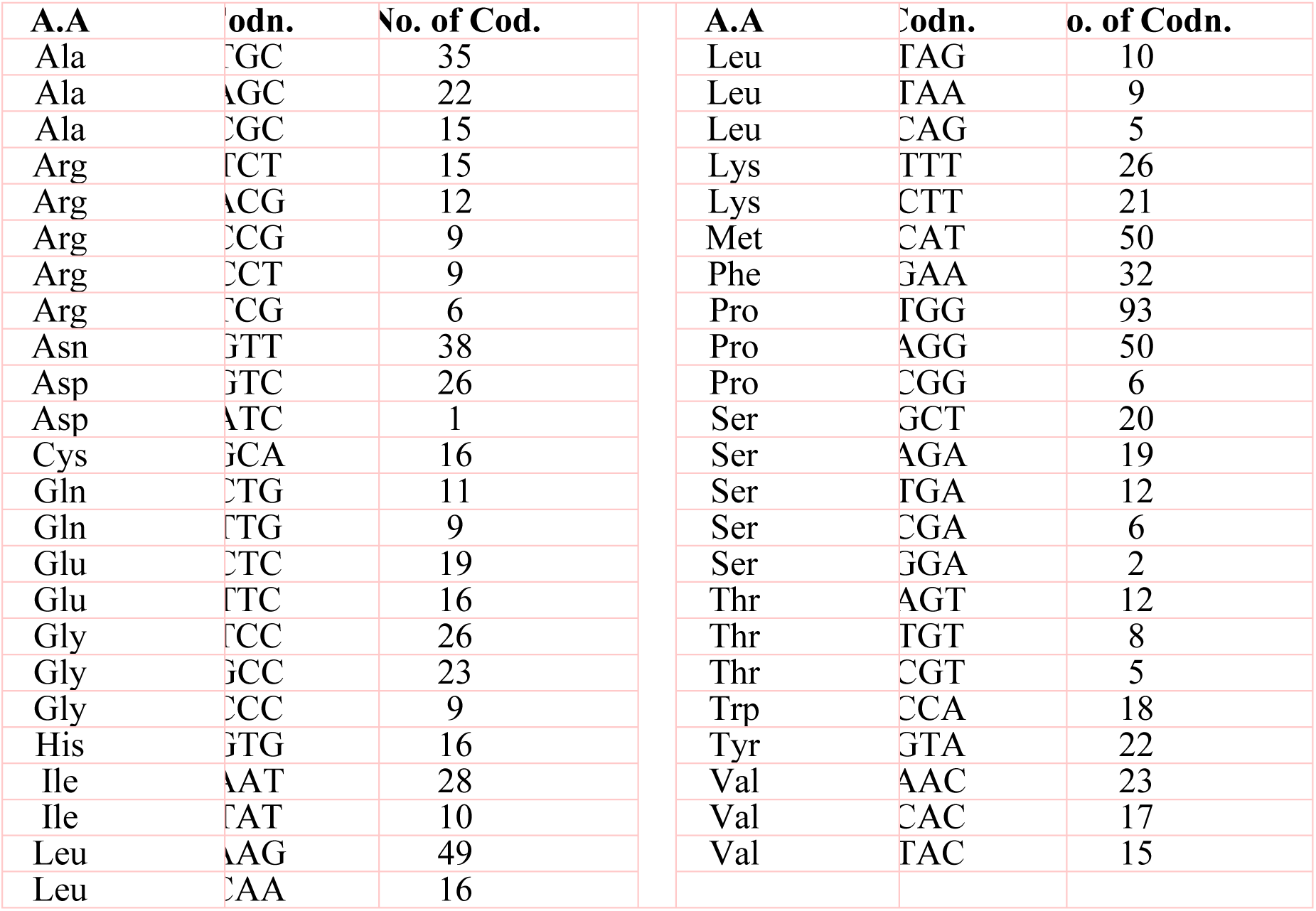
tRNA prediction response to aminoacid combinations and codon distributions

A total of 34 tRNA bound regions were found in intronic spacers, with internal scores between 92.4-84.2. RNA self-splicing is predetermined by these regions, which tends to translate the number of amino acids from the predicted genes (Smathers and Robart, 2019). A total of 543 scaffolds contributed to bound tRNA genes, as indicated in Table 10. A total of 577 tRNA loci (excluding 34 pseudo tRNAs) spanning 430 bp with an average length of 20 bp were predicted.

Interestingly, only tRNA genes encoding Tyr, Met, Lys, Asn, Ser, Ala and Leu showed the presence of introns. The average intron length of the tRNA genes for Tyr and Met was 20 bp, while for Lys, Asn, Ser, Ala and Leu, these lengths were 19 bp, 14, 10, 6 and 4 bp, respectively. The predicted tRNA genes indicated the maximum density of copies on the scale of the assembly contribution.

#### rRNA cluster fragment distribution

The rRNA genes in the assembly were calculated with RNAmmer 1.2. Only full model hits with E-values <0.01 were accepted as reliable hits, and most of the hits with E-values between 0.01 and 1 were reported. The alignment for rRNA prediction was placed farther downstream to increase the good number of model scores. Twelve full-length tandem arrays (large and small units) of rRNA genes were found in the assembled scaffolds (Fig 6). In total, 4 scaffolds were involved in the rRNA prediction calculation, and we observed an average length of 672 bp. The highest match of 4568 bp of rRNA was found in scaffold number 8716. 8S, 18S and 28S rRNA units were predicted to have lengths of 113, 1806 and 4586 bp, respectively. One positive strand was related to 8S rRNA genes of approximately 114 bp length. Only full model hits with E-values <0.01 were accepted as reliable hits, and no hits with E-values between 0.01 and 1 were reported (Table 11). The variable scores of the unique rRNA operon predictions did not significantly differ from the annotated values, indicating that true rRNAs had not been annotated from the GenBank annotations. No positional overlap was found between the predicted tRNA and rRNA scaffold regions.

**Fig 6:**
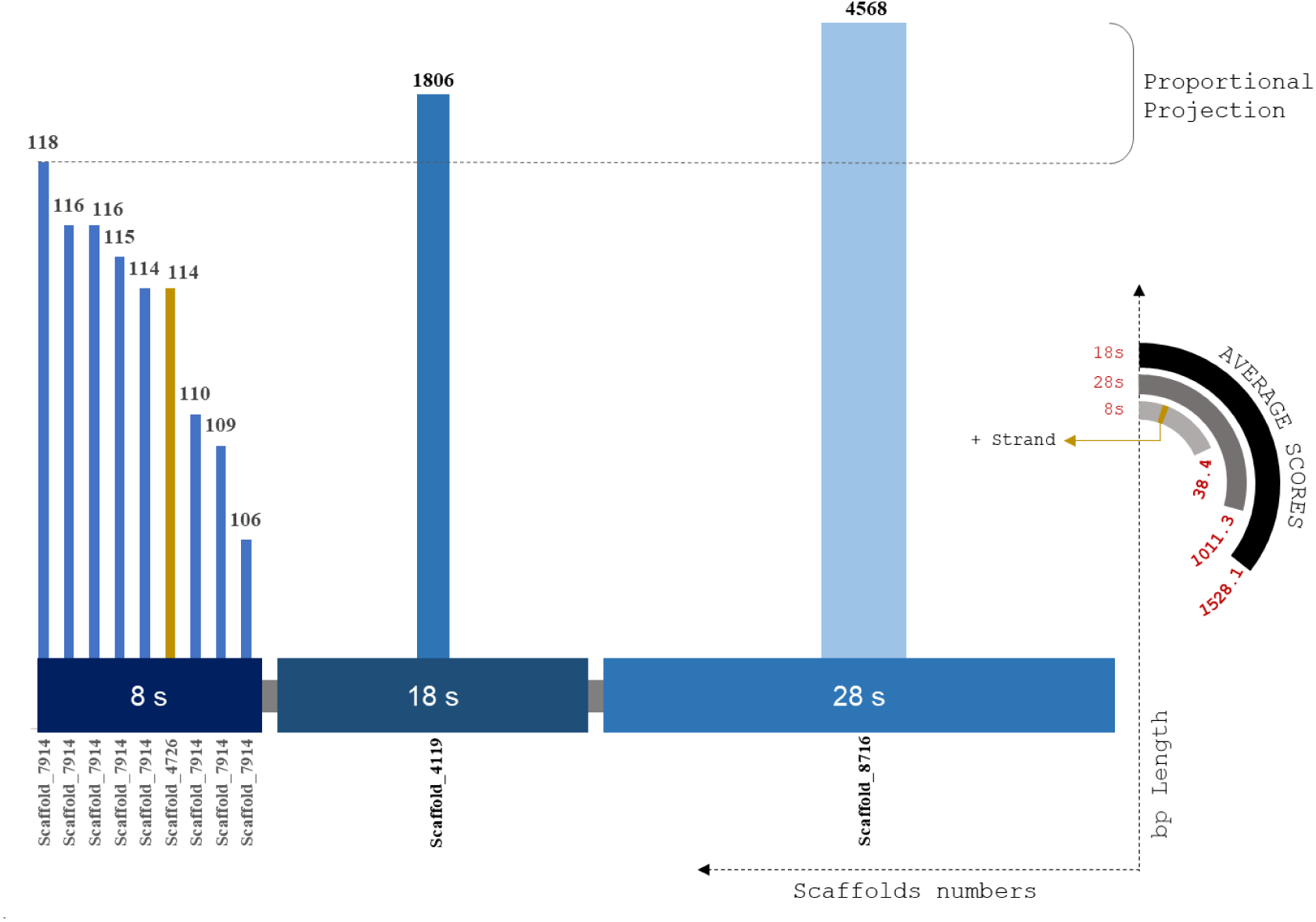
rRNA plot by distributed size of ribosomal RNA subunits against the scaffold utilised for prediction using RNAmmer. X coordinates are subunit classification and y coordinate classifies scaffold count and length of each scaffolds. The pseudo semi circular projections depicts the score value of each prediction and strand profile of predicted rRNA profiles.

**Table 11:**
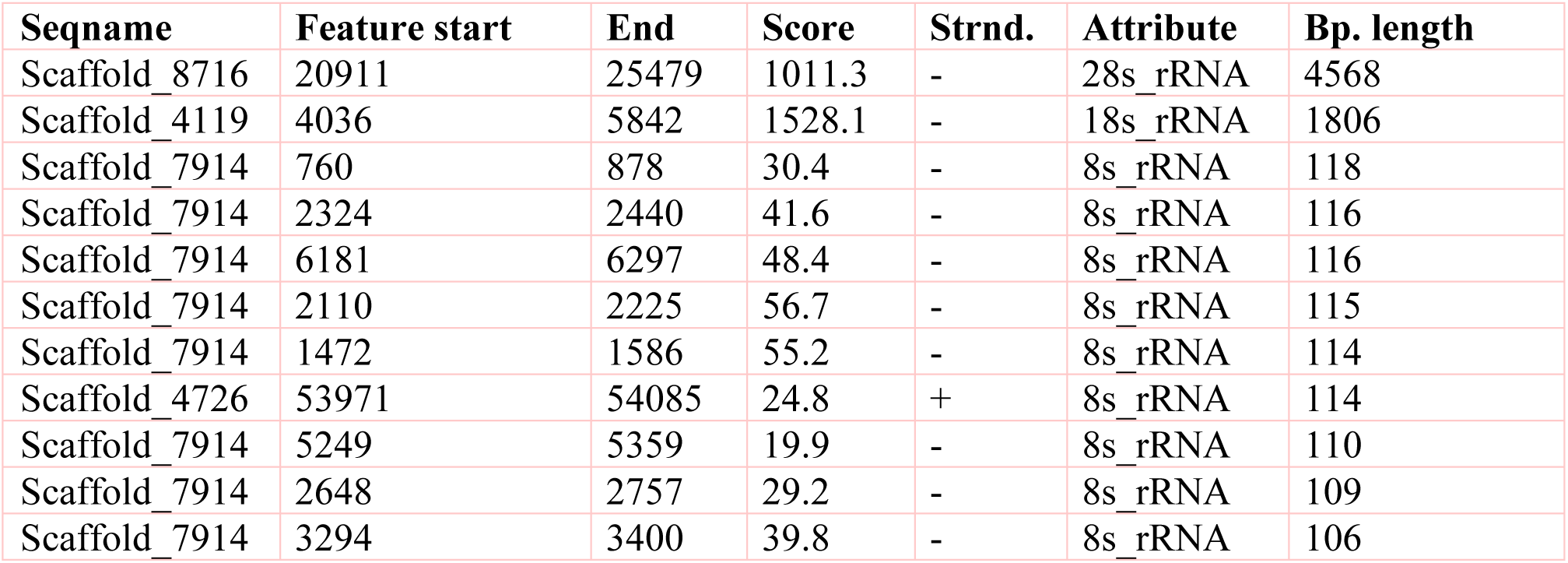
Complete rRNA sequence predicted from scaffolds assembled raw reads. 4 scaffolds (Scaffold 8716, 4119, 7914 and 4726) are involved in the prediction of 11 rRNA sequence for the distribution of 28s, 18s, 8s subunits.

### SSR marker analysis

***Compound Repeat Annotations – Class I:*** Nucleotide diversity analysis of *Vigna umbellata* draft assembly provided a rich resource of polymorphic SSRs, which are useful to determine marker-trait relationships, especially for quantitative trait loci and molecular breeding. Repeated operons for hundreds of polymorphic non-monomeric SSRs and flanking bases from the assembled data were catalogued to translated them into genetic markers (Vining *et al.,* 2017). The repeats were categorized based on the position and number of repeats of each allele. The compound repeats (Cr) were calculated for stable (c) and variable (c*) alleles (Supplementary table - 2). C_r_ presented a maximum scaffold length of 9711 bp for the stable allele (SA), while the variable allele (VA) presented a length of 1048 bp. The length of the repeat loci was 71 bp on average (median statistics) for SA and 92 bp for VA. The number of C_r_ poly-hits ranged from 2x – 54x, whereas the 32 single ranged from 19-5284 combination repeats (Fig 7).

**Fig 7:**
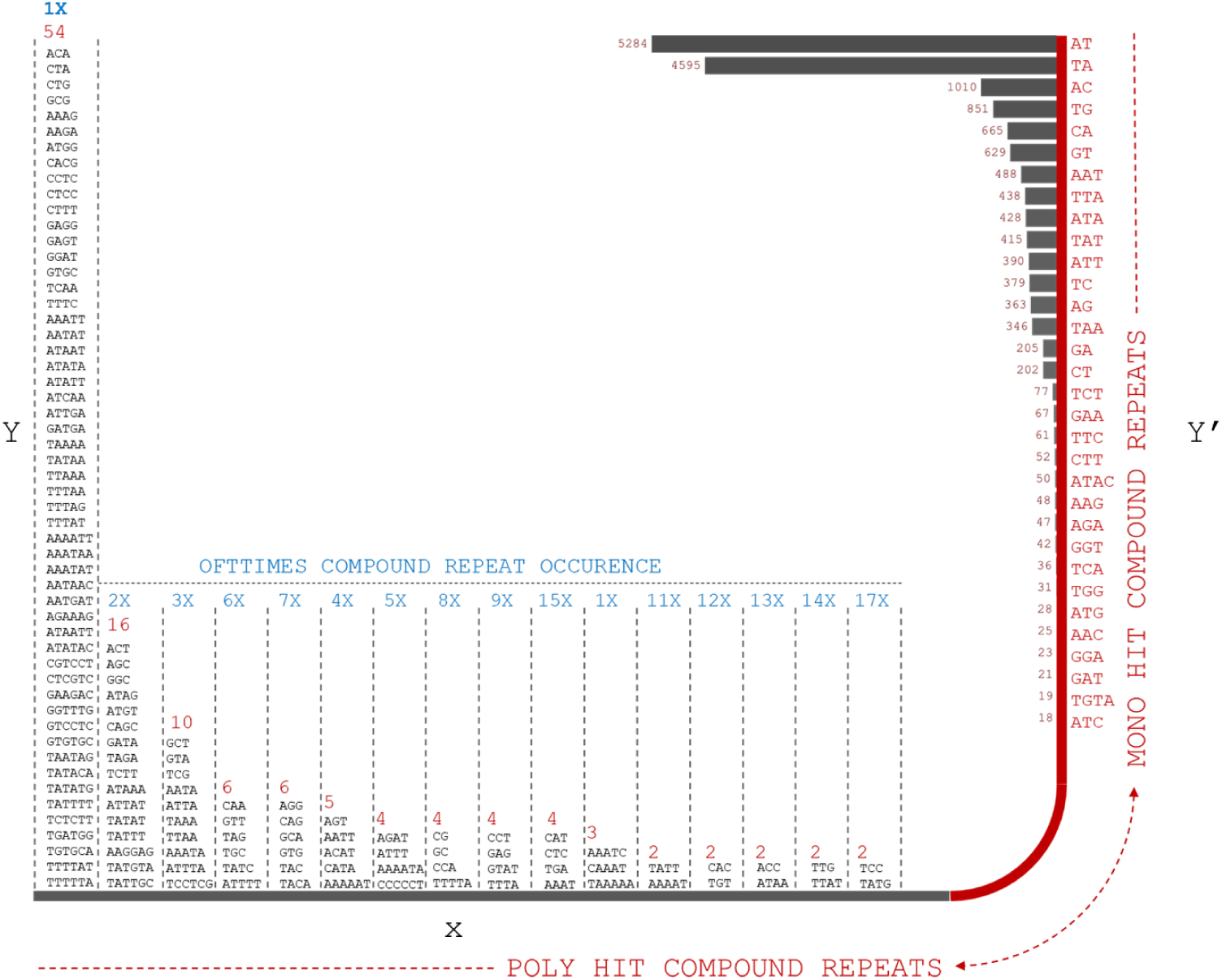
Compound repeats from assembled reads were plotted from the distribution of offtimes of repeat occurrence and number of hit (mono and poly hits). X-Y coordinate prevails the distribution of compound repeats in multiple combinations whereas X-Y’ projects mono repeats with multiple form bp combinations.

#### Stemplot distribution of non-Compound SSR markers [P2,P3,P4,P5,P6]

**Hexa-nt repeats** (Fig 8): The stemplot was computed for non-compound SSR repeats. A total of 2111 (0.1%) six-nucleotide repeats were analysed. The calculations were performed to understand the number of continuous bp in each P2 repeat. The frequency distribution of repeats varied in the series of 5x[A:T], 4x[A:T:G:C], 3x[A:T:G:C], 2x[A:T:G:C] and Random. The maximum number of distribution series was 24 x 2CX [24 combinations of X 2 ‘C’ repeats in the SSR; for example: AAGA**<u>CC</u>**], in which the lowest was 3 x 5AX. **Penta-nt repeats** (Fig 9): a total of 2045 (0.1%) P5 repeats were predicted from the assembled scaffolds. The frequency distribution of the repeats varied in the series of 4xA:T:C, 3xA:T:G:C, 2xA:T:G:C and Random. The maximum number of distribution series was 24 x 2CX, in which the lowest was 3 x 5AX. **Tetra-nt repeats** (Fig 10): 29524 (1.8%) were found in continuous four-nucleotide base repeats - 3x[A:T:G:C], 2x[A:T:G:C] and Random. The maximum frequencies showed random combinations, and the lowest presented 3xG and 3xC. **Tri-nt repeats** (Fig 11): 76493 (6%) 3 bp repeats were found, with a maximum frequency of 36 from random and the lowest with all 2x A:T:G:C leaflets. **Di-nt repeat** (Fig 11): 1297040 (92%) random combinations of 2 bp repeats were identified from the assembled reads.

**Fig 8:**
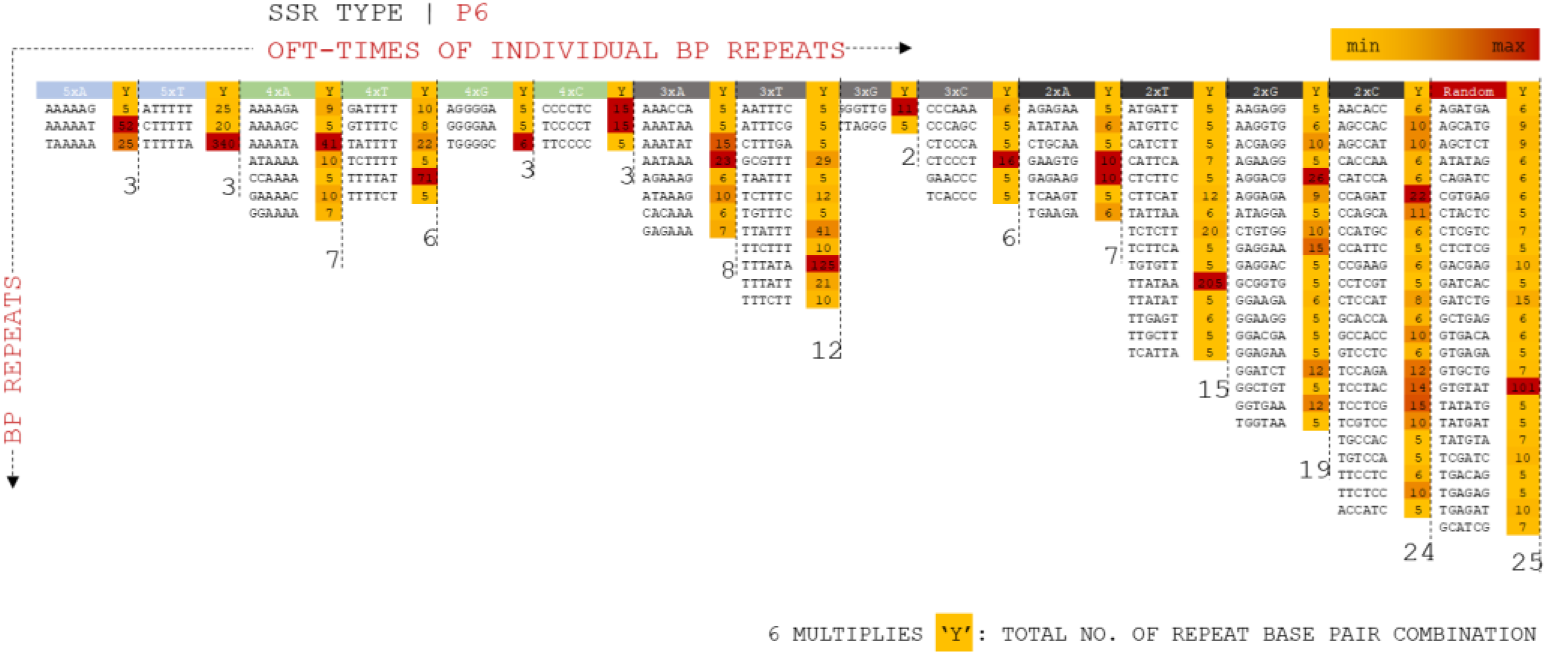
P6 SSR operons mapped for distributed combinations for 5xX (A, T, G, C), 4xX, 3xX, 2xX and random specificity of repeats. Operons are projected on positioning oft-time count, Y mulitiples of each marker types.

**Fig 9:**
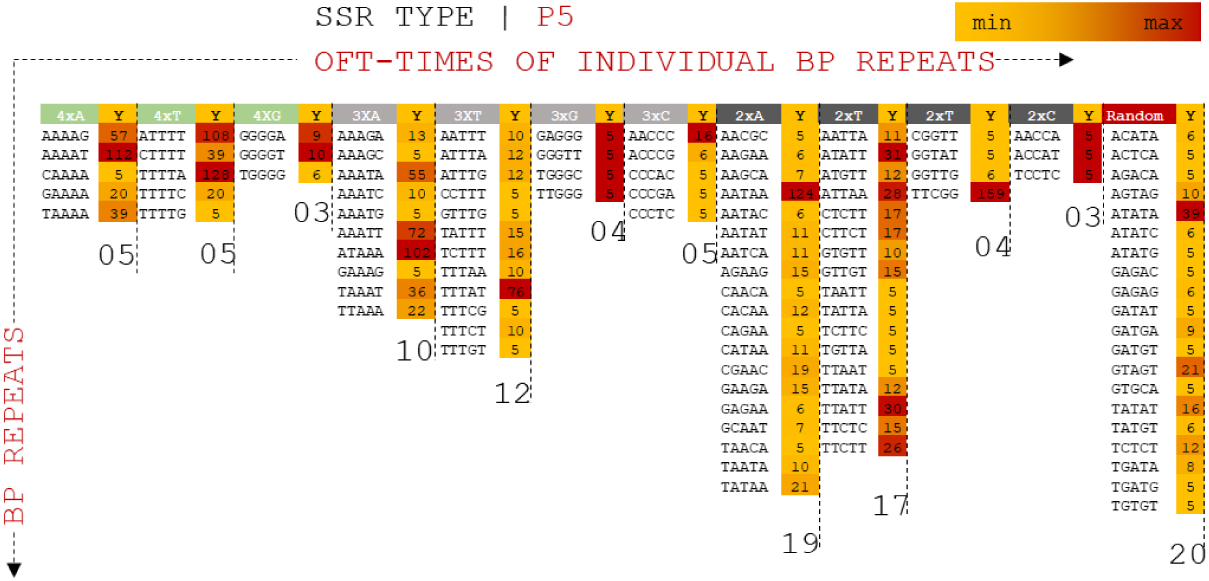
P5 SSR operons mapped for distributed combinations for 4xX(A, T, G, C), 3xX, 2xX and random specificity of repeats.

**Fig 10:**
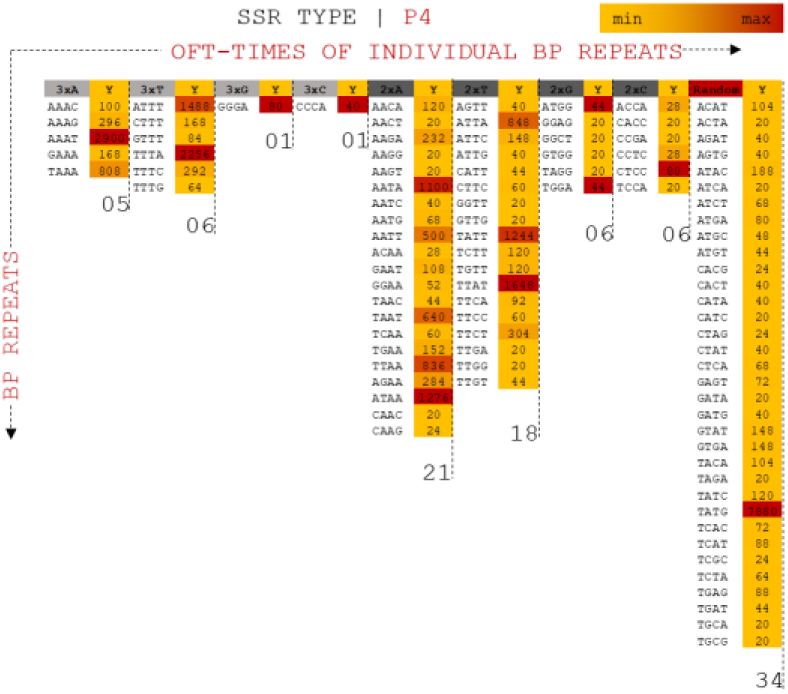
P3 SSR operons mapped for distributed combinations for 3xX(A, T, G, C), 2xX and random specificity of repeats.

**Fig 11.**
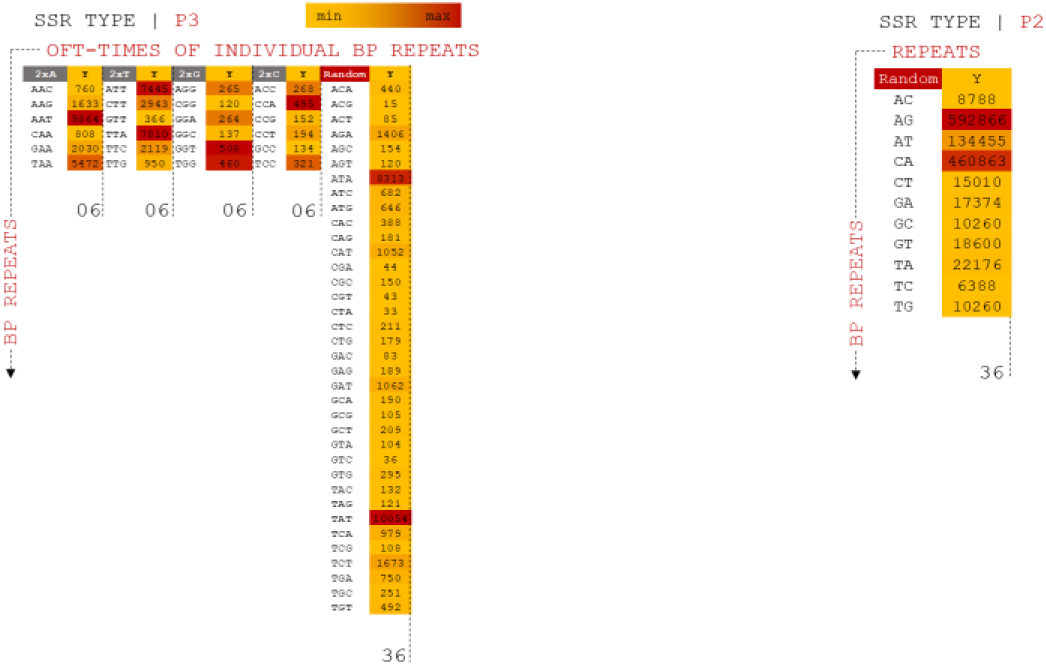
(a) P3 SSR operons mapped for distributed combinations for 2xX(A, T, G, C) and random specificity of repeats. (b) P2 SSR operons mapped for distributed combinations for random specificity of repeats.

#### BLASTx Analysis

All predicted genes were annotated by evaluating their homology through BLASTX searches against the NR database (non-redundant database) for complete integration with the physical/genetic maps (Zhao *et al.,* 2019). The genes were aligned using the following optimized search parameters: gap opening penalty (G) = 4, gap extension penalty (E) = 1, mismatch score (q) = −1, match score (r) = 1, word size (W) = 11 and e-value <e−20 (Singh *et al.* 2004). Low-complexity filters were enabled in the total search parameters. Based on similarity searches against known proteins and genes, with a cut-off E-value of 10, the total number of genes predicted from the assembly was 31276, including 31237 genes with BLAST hits and 39 genes with no BLAST hits. The average gene length was 1.8 kb, and the average GC content was 42%. The maximum length was 12091 bp for scaffold 2254 (serine threonine-kinase SMG1-like [XP_017432882] from *Vigna angularis*) with approximately 99.58% similarity. A total of 7708 scaffolds were associated with enzymes: 279 were membrane specific; 1418 were related to the chloroplast response; 759 were related to the mitochondrion response; 442 were ribosome specific; 1082 were transcription related; 89 were related to leucine rich-repeat regions and 69 were translational related. *Scaffold percentage distributions:* The top species hits ranked 1 for *Vigna angularis*, secondly with *Vigna radiata* and *Vigna unguiculata* as third closer genome, which were calculated from the assembled reads. The total % distribution ranged from 100 – 43. Major crops such as *Glycine max*, *Glycine soja*, *Zea mays, Vigna aconitifolia* and the *Oryza sativa* japonica group presented a throughout the alignment frequency. A total of 90% of the crops from the 100% index fell within the 99-97% index matches. The counts of scaffolds are proposed to be used to study the number of scaffolds (Fig 12.a). A total of 6235 scaffolds came from the 100% distributions, 2139 scaffolds from 99-90%, 2297 scaffolds from 89-80%, 971 scaffolds from 79-70%, 284 scaffolds from 69-60%, 47 scaffolds from 59-51%, and 10 scaffolds from 49-43%. *Species Distribution from the % index:* The distribution of species other than crops screened from the 100% index was evaluated. The total frequencies of the species distributions were 44% from crops, 15% from animals, 10% from fruit, 9% from flowers and trees, 3% from birds and vegetables, 2% from insects and 1% from bacteria and coral (Fig 12.c). Divergence plots are classified based on crops and maximum hits. A total of 25771 scaffolds came from *Vigna angularis*, 4840 from *Vigna radiata*, 223 from *Phaseolus vulgaris*, 88 from *Glycine max*, 78 from *Cajanus cajan*, 22 from *Glycine soja*, 15 from *Trifolium pratense* and 22 from other species (Fig 12.b). The total alignment percentages, scores and functional annotations of each gene from the corresponding species are listed in Supplementary Table 3.

**Fig 12:**
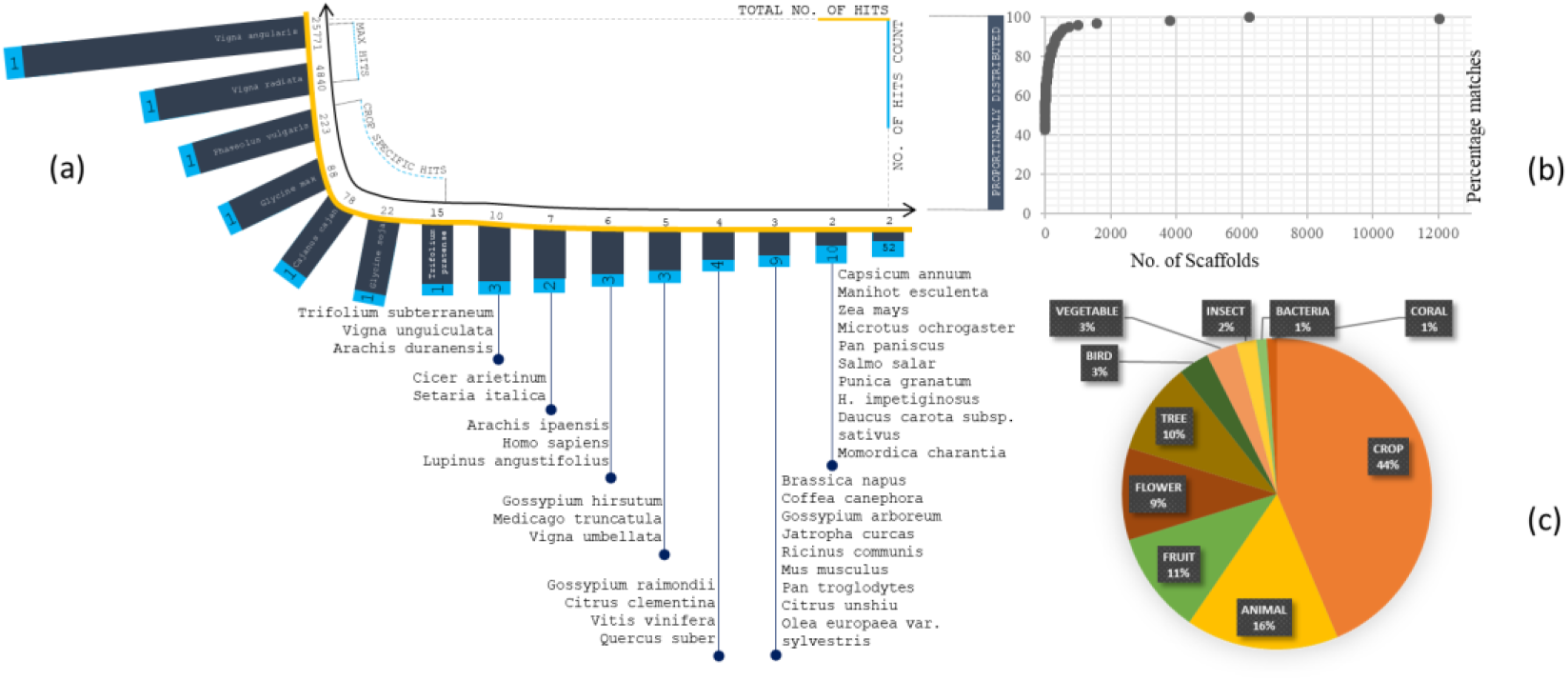
(a) Distribution of Species from total scaffolds matched from blastX Analysis. Coordinates projected in terms of scaffold matches (total no. of hits and no. of hits counts) (b) Percentage distribution of scaffolds in correspondence of species fall. (c) Plot on total organism matches with its generic names.

#### GO analysis

GO sequence distributions help to identify the annotated nodes comprising GO functional groups. Functional descriptions were assigned to predicted *Vu* genes. Genes associated with similar functions are assigned to a similar GO functional group (Xu *et al.,* 2017). This study identified several potential genes involved in the response to palatable indexed genes, flowering potential, stress response, and disease resistance genes to various metabolic processes and the regulation of transcription and transport. The total number of GO-annotated genes was 16974, among which the overlapping positional map included 11204 biological processes, 13097 molecular functions and 9053 cellular components predicted from the genes. These findings should help to identify and manipulate *Vu* genomic diversity to contribute to the development of crops.

*Biological Process:* A total of 166622 independent annotations with 23907 dependent genes were assigned GO terms, among which the genes predicted from the assembly exhibited 11204 functions related to biological processes. The 6957 prevalent functions corresponding to Biological Processes from ricebean were related to ion-dependent functions, with predominance of membrane-transporter activity and haemostasis. The second largest category among the genes associated with DNA dependence included approximately 1560 functions involved in replication, transcription, translation, endonucleolysis, repair mechanisms, topological changes, duplex unwinding and signal transduction. The approximately 918 identified membrane functions play critical roles in transport mechanisms such as ion, electron, antigen, amino acid and ATP transport as well as signalling and response mechanisms. Major notable biological processes were assigned to top functions such as the following categories: GO:0008152-[8670; scaffold fragments] metabolic process, GO:0009987-[7253] cellular process, GO:0065007-[2401] biological regulation, GO:0051179-[1550] localization, GO:0050896-[1229] response to stimulus, GO:0071840-[936] cellular component organization or biogenesis, GO:0023052-[478] signalling, GO:0032502-[272] developmental process, GO:0032501-[272] multicellular organismal process, GO:0000003-[182] reproduction, GO:0051704-[142] multi-organism process, GO:0002376-[43] immune system process, GO:0040007-[34] growth, GO:0048511-[14] rhythmic process, GO:0015976-[6] carbon utilization, GO:0040011-[6] locomotion, GO:0008283-[5] cell proliferation, GO:0022610-[5] biological adhesion. The detailed and expanded version of the biological process results was plotted as shown in Fig 13. *Cellular component:*A number of scaffolds were associated with cellular components, and the corresponding functions were predicted for the mapped genes. A total of 17 cellular components were reported from the prediction analysis. A total of 5347 components were identified from membrane-specific components in the plot maximum, combining five functional sub-components. The membrane components were further subjected to location identification based on their functions (e.g., cell wall, chloroplast, endoplasmic reticulum, Golgi membrane or vasculature) (Fig 14). A total of 1534 components identified from the nucleus corresponded to 38 co-functional components. All other cellular components with multiple subcategories, including the Cytoplasm, Chloroplast, Ribosome, Intracellular, Golgi, Mitochondria, Thylakoid, Photosystem, Chromosome, Enzyme, Plasmodesma, Endosome, Microtubule, Vacuole and Endoplasmic reticulum were distributed according to their locations as follows: 348 x 11 counts, 315×27, 245×19, 239×38, 167×87, 125×6, 92×19, 68×12, 38×8, 38×8, 33×42, 26×17, 15×13, 12×223 and 5×70, respectively. *Enzyme Functions:* The GO-profiled enzymes were classified into seven categories: oxidoreductases, transferases, hydrolases, lyases, isomerases, and ligases, which varied depending on the responses that the enzymes catalysed (Morgat *et al.,* 2019). The most common types of enzymes were oxidoreductases, transferases and hydrolases. Distinct enzyme classes were further systematically categorized based on the substrate reaction and mechanism of response. The composite enzyme contributions were categorized as E.C 1-2-3-4-5, while there were a total of 184 top-level classifications and 10531 complete classifications (Fig 15). The highest distributions were accompanied by approximately 1464 major | 64 subclassified E.C 1.1 reaction types, e.g., electron transfer: A^+3^+B^+2^- >A^+2^+B^+3^. Following the electron transfer mechanism, 311 | 58 enzymes were classified as E.C 2.1; state of functions derived from the transfer of functional groups; the reaction type is A–X - > A+B-X. The total number of enzymes from the 3.1 nomenclature was 282 | 41 predicted; the reaction type was hydrolysis [A-B+H_2_ 0 - > A-OH + B – H]. E.C 4.1 and 5.1 are 1 | 5 and 11 | 2 combinations of reaction types, respectively (Table 12); i.e., elimination to form double bonds [(A-X)-(B-Y)->A=BX-Y] and transfer of groups within molecules [(A-X)-(B-Y)- >(A-Y)-(B-X]. The complete, curated enzyme classifications are provided in the supplementary table 3. *Other Molecular Functions:* In addition to enzyme functions, four other functions were predicted from the GO analysis: Binding proteins 194 distributions in 4189 genes, Transporter 38 distributions in 522 genes, Channel 13 distributions in 87 genes, Dogmatic 7 distributions in 88 genes and Inhibition/Activation-specific 40 distributions in 855 genes. There were 292 | 5741 genes major and sub-classified functions in the cellular mechanism category (Table 12).

Fig 13: Biological Process annotated from GO profile. Totally 45 BP distributions analysed from assembly.

**Table 12:** Functions of genes mapped from rice bean distributed for its scaffold count and total number of count. Complete distributions are concerning enzyme classifications and other functional activity of the mapped genes.

| Functions | Fragment count | Total count |
| --- | --- | --- |
| EC.1_Oxidoreductases | 64 | 8550 |
| EC.2_Transferases | 59 | 1017 |
| EC.3_Hydrolases | 41 | 683 |
| EC.4_Lyases | 5 | 8 |
| EC.5_Isomerases | 2 | 19 |
| Binding | 194 | 4189 |
| Transporter | 38 | 522 |
| Channel | 13 | 87 |
| Dogmatic | 7 | 88 |
| Inhibition and activation | 40 | 855 |

**Fig 14:**
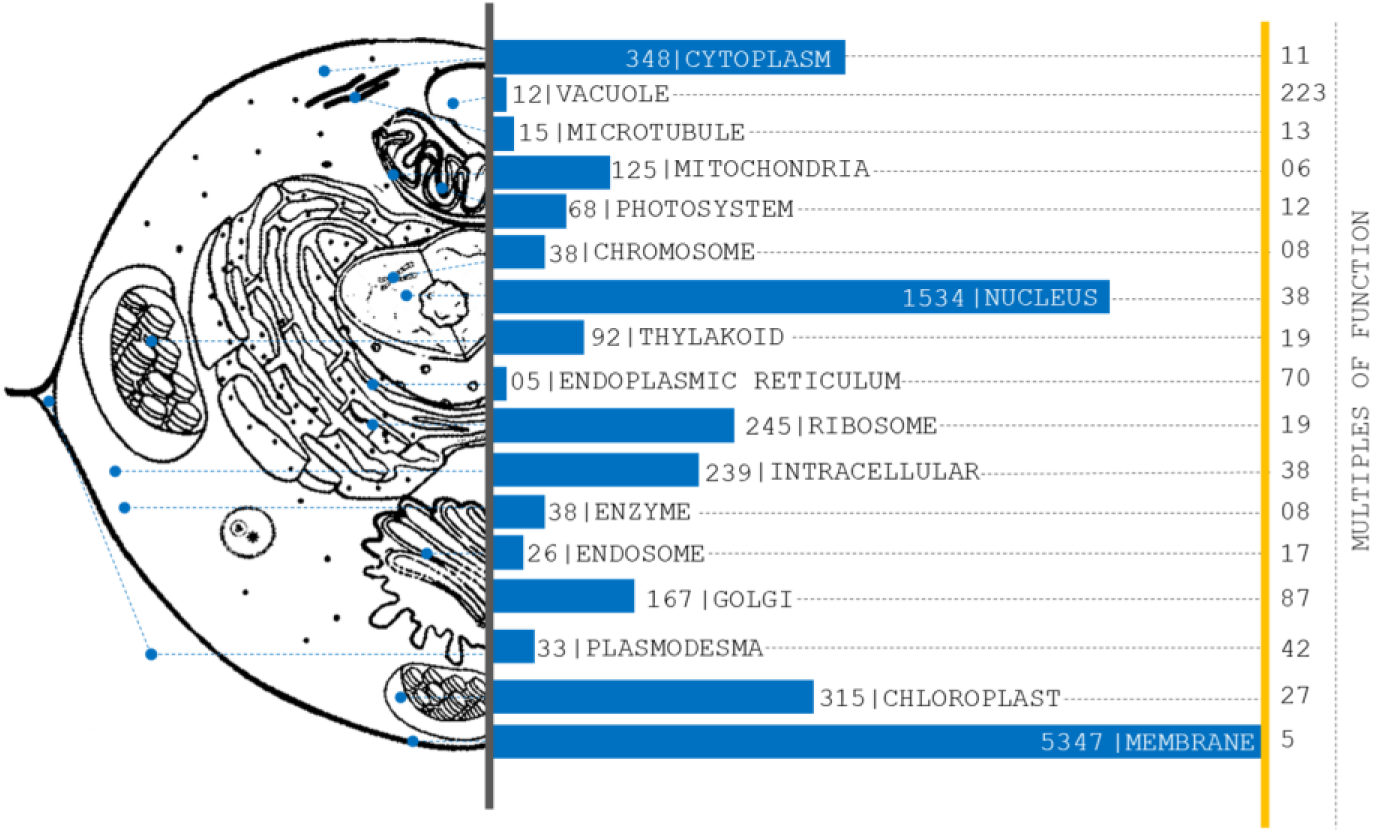
Localisation of Organelle of scaffolds from assembly predicted by GO profile. Totally 555 leaflets of organelle found and categorized to plot for 17 various major organelles in plant cells. Histogram plotted proportional to number of counts per organelles from GO predictions.

**Fig 15:**
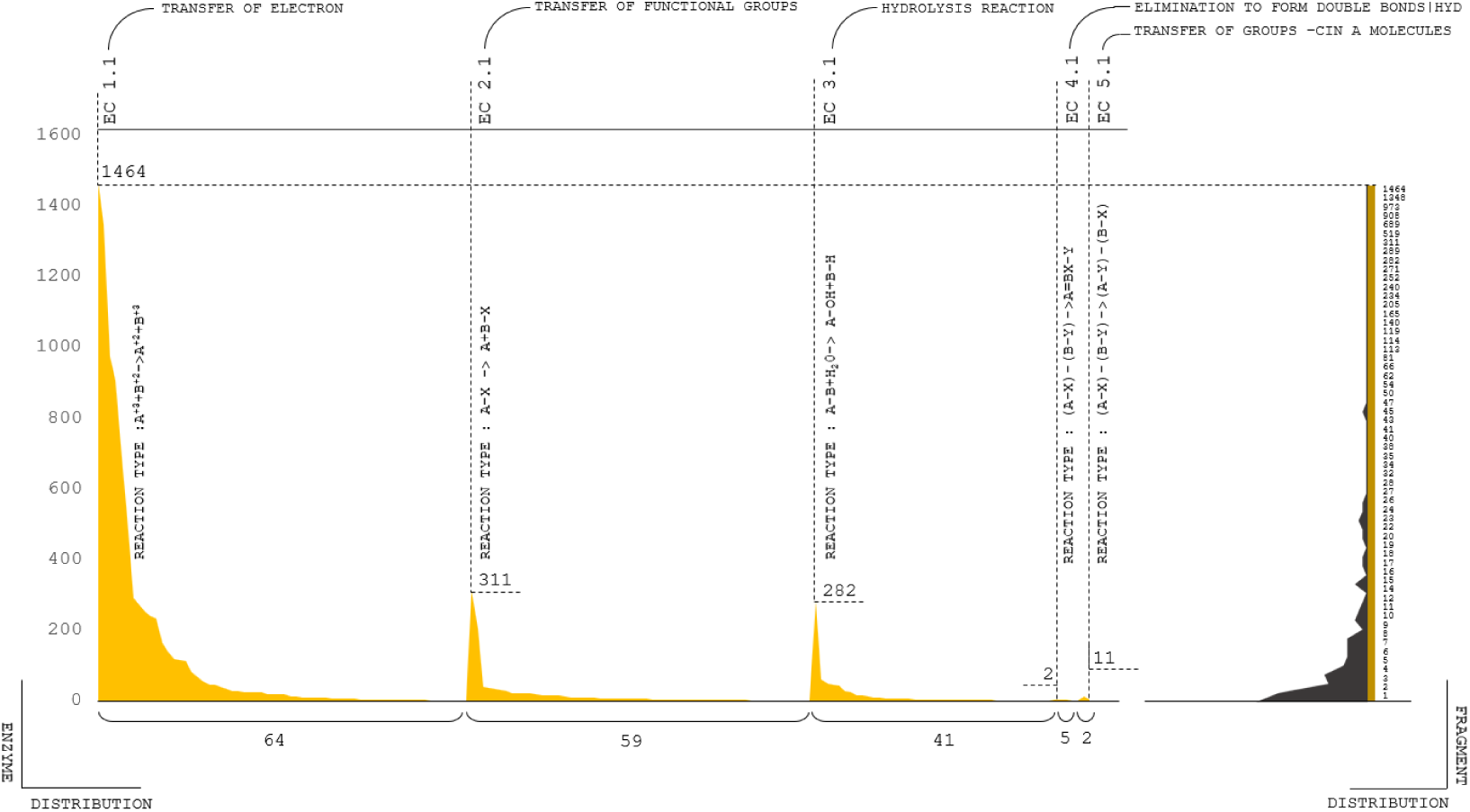
Enzyme composition predicted from scaffolds and projected the plot based on enzyme classification and enzyme reactions types. Two coordinates corresponds the scaffold distribution and no. of enzyme reaction types. Totally five enzyme classifications were analysed and plotted against reaction types and scaffold distributions.

### KEGG orthologue assessment

A total of 515 functional orthologs were derived from 19173 experimentally characterized KEGG-gene associated pathways. The pathway annotations included the functional hierarchies and binary relationships of biological entities.

#### Gene distribution in KEGG annotation pairs

The 54 functional hierarchies were predominantly related to the correspondence of orthology with KO identifiers (Ho, Lee and Lim, 2018). Twelve metabolic hierarchies included 167 orthologs. Four genetic information processing hierarchies exhibited 22 orthologs. Three environmental information processing hierarchies exhibited 40 orthologs. Five cellular process hierarchies exhibited 33 orthologs. Ten Organismal system hierarchies exhibited 87 orthologs. Twelve human disease hierarchies exhibited 89 orthologs, and 4 BRITE hierarchies exhibited 53 orthologs (Fig 16). The significance of the composite gene distributions for the hierarchies demonstrated functional consistency in the metabolic resources and disease-specific mechanisms. Triangulation of multiple vertices of pathway annotations empirically contributes to the expectancy of obtaining ricebean genome annotations. ***Genes with orthology pairs:*** Calculation of the number of predicted rice bean genes involved in the KO and hierarchy predictions was performed to understand the functional dynamics of *Vigna umbellate* (Fig 17). The metabolism-specific orthologues primarily indicated the functions derived from KO prediction. The metabolic pathways for the percentile plot were as follows: 9% (15 reference hierarchies and 3595 pathway hits) Carbohydrate metabolism, 5% (8 reference hierarchies and 377 pathway hits) Energy metabolism, 10% (17 rh and 442 ph) Lipids, 1% (2 rh and 156 ph) Nucleotides, 12% (23 rh and 714 ph) f Amino acids, 11% (18 rh and 160 ph) Glycan biosynthesis, 7% (12 rh and 241 ph) cofactors and vitamins, 13% (21 rh and 154 ph) terpenoids and polyketides, 17% (29 rh and 247 ph) Biosynthesis of other secondary metabolites, and 13% (21 rh and 150 ph) Xenobiotic biodegradation. The pathways related to genetic information processing were attributed to for four primary mechanisms: 13 % (3 rh and 267 ph) Transcription, 23% (5 rh and 643 ph) Translation Folding, 32% (7 rh and 470 ph) sorting and degradation and 32% (7 rh and 268 ph) Replication and repair. The environmental information processing category included 7% (3 rh and 27 ph) Membrane transport, 83% (33 rh and 1258 ph) Signal transduction and 10% (4 rh and 0 ph) Signalling molecules and interactions. The cellular process category included 27% (9 rh and 581 ph) Transport and catabolism, 37% (12 rh and 541 ph) Cell growth and death, 15% (5 rh and 121 ph) Cellular community - eukaryotes, 12% (4 rh and 73 ph) Cellular community - prokaryotes, and 9% (3 rh and 66 ph) Cell Motility. BRITE hierarchies are resources for understanding the high-level functions and utility of biological systems (Kanehisa and Sato, 2019). According to these hierarchies, 25% (13 rh and 1211 ph) were assigned to Protein families-metabolism, 30% (16 rh and 4575 ph) to Protein families-genetic information processing, 43% (23 rh and 1212 ph) to Protein families-signalling and cellular processes, and 2% (1 rh and 0 ph) to the RNA family. There were 386 poorly characterized orthologs, among which 4 reference hierarchies and 24 pathways were studied (Fig 18).

**Fig 16:**
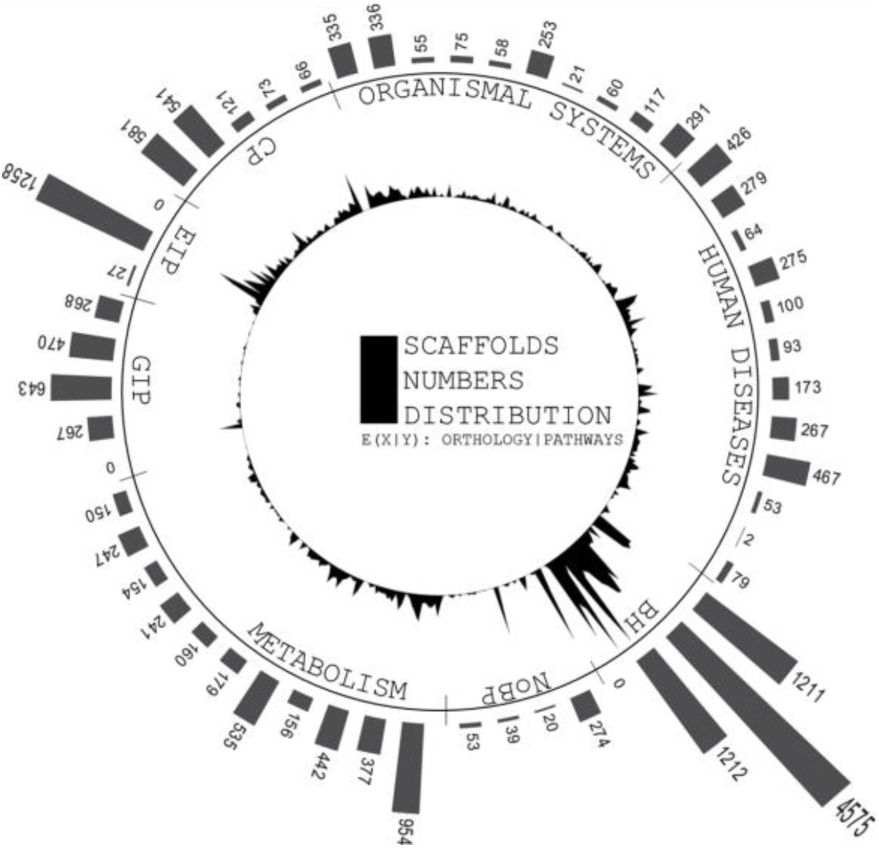
Circular and distribution peaks are projected from number of scaffolds involved in each orthologs. First layer of circular projections shows the maximum peaks of scaffolds distributions from each pathway categorized in orthologs. Second layer depicts the number of pathways with respect to scaffolds involvement to predict its pathway functions.

**Fig 17:**
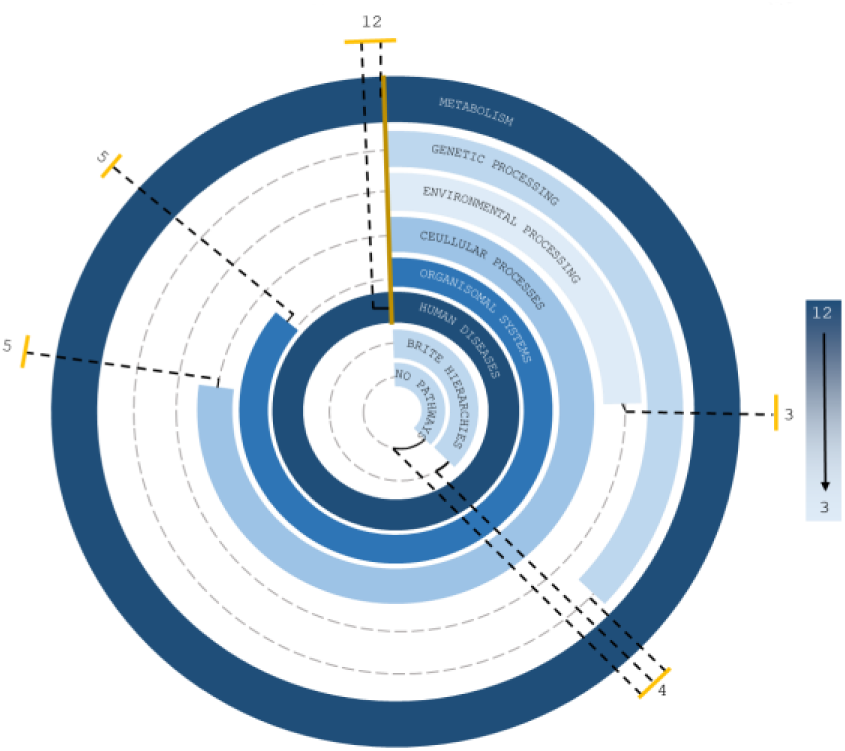
Circular projections of Pathways hits from assembled scaffolds of Rice bean. The plot coordinates are differentiated between number of orthologs and total number overhead pathway counts from each prthologs from complete prediction profile.

**Fig 18:**
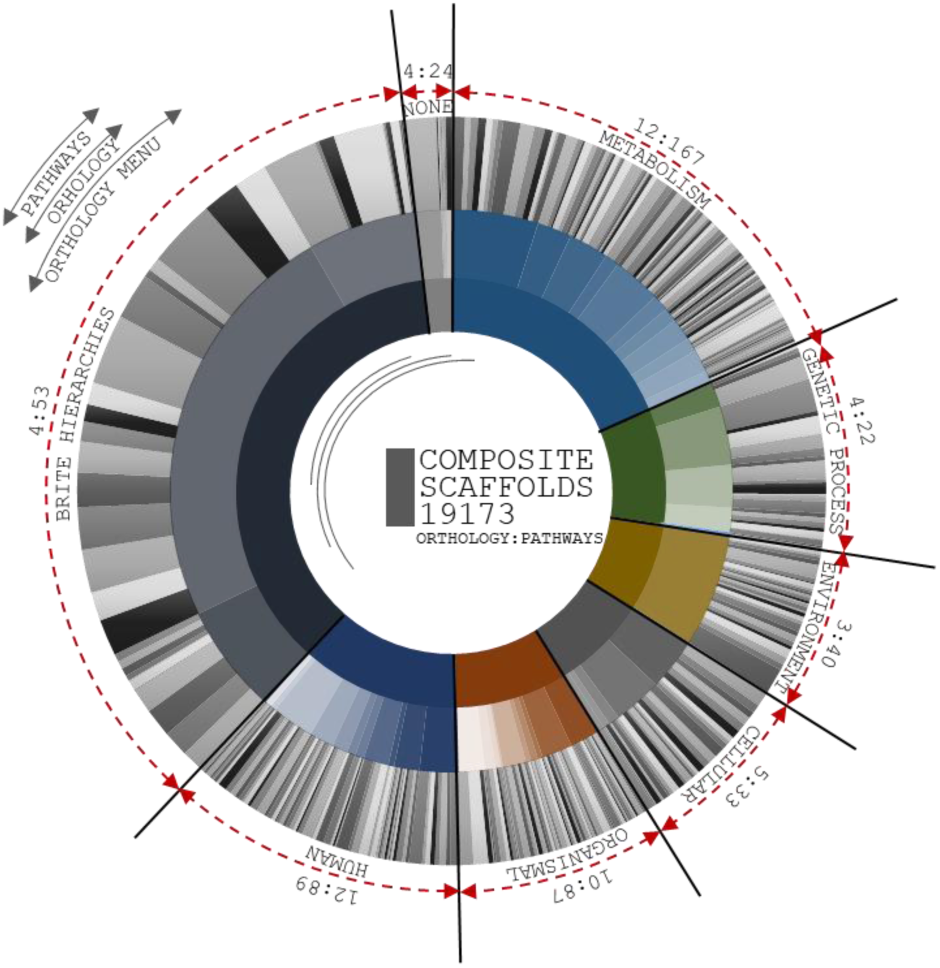
Circular distributions of pathways from ortholog head. Composition of three layers are coordinated by number orthologs, number of pathways from each orthologs, and scaffolds distributions at each pathways. First Layer projected for number of orthologs from KAAS server, second layer depicts the number of pathways from seven ortholog head and third layer distributed based on scaffold involved in each pathways predicted

#### Functional distribution of ortholog pairs

Functional hierarchies were plotted on the basis of orthology to KO distributions (Fig 19). The greatest number of functional pathways hits were for protein families such as those involved in genetic information processing (4575 pathway hits) and signalling and cellular processes (1212 pathways). In the total orthology count retrieved from the KASS server, 3% of GIP included 13% Membrane trafficking and 12% chromosomal and associated proteins. Four percent of signalling and cellular processes included 84% of Exosomes and 38% transporters in the pathway predictions. The assessment of complete pathway patterns revealed that 7% of Signal transduction included 17% Plant hormone signal transduction and 6% Protein families; metabolism included 18% Protein kinases, 18% Protein phosphatases and associated proteins, 19% Peptidases and 17% Glycosyltransferases. Among 24% of Protein families, the composition of the genetic information processing category was as follows: 8% Ubiquitin system, 8% Messenger RNA biogenesis, 8% Spliceosome, 7% DNA repair and recombination proteins, 6% Transcription factors, 6% Ribosome, 6% Ribosome biogenesis and 5% Transcription machinery. In the 1% of Unclassified metabolism, 91% consisted of Enzymes with EC numbers.

**Fig 19:**
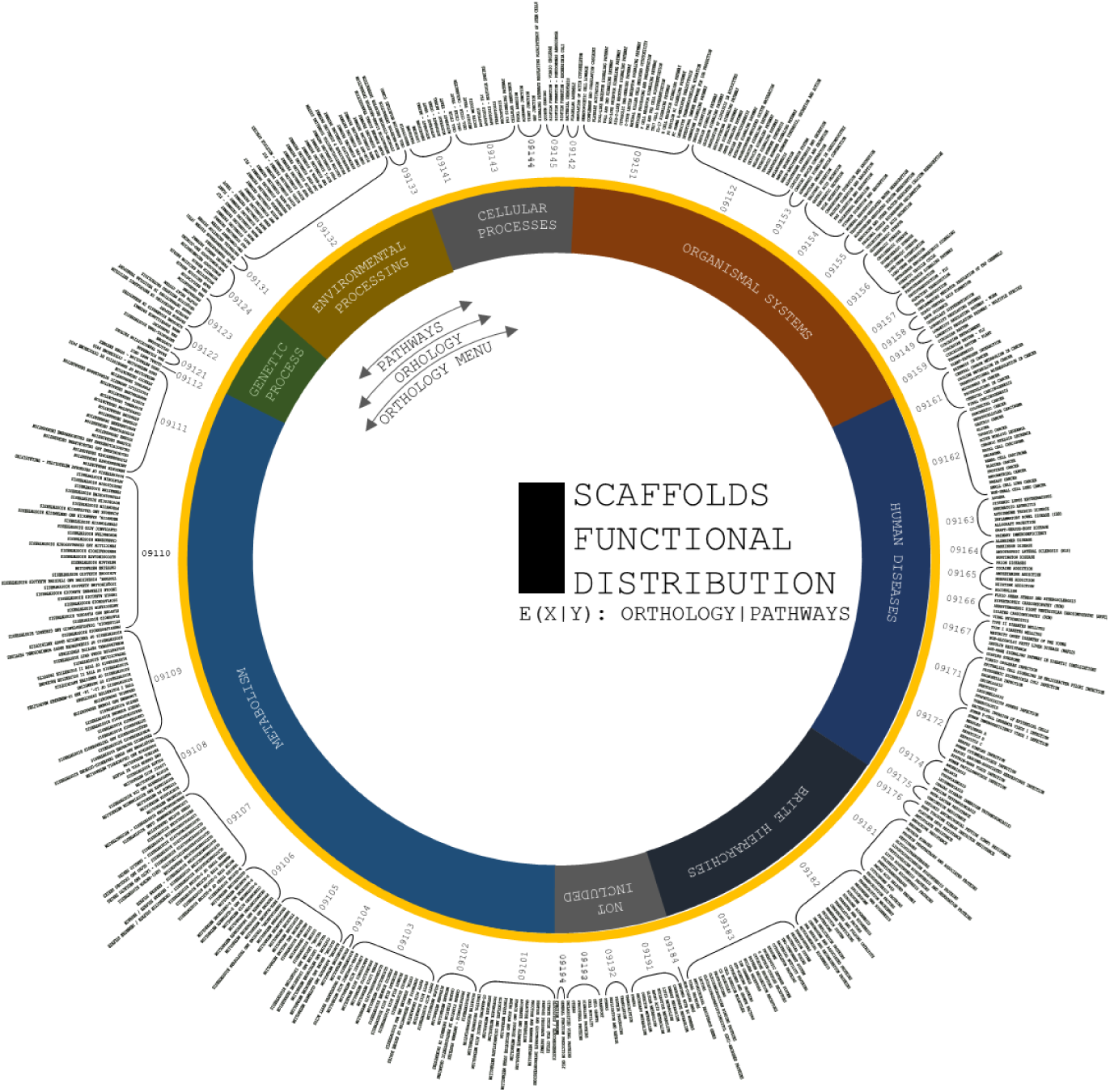
Circular projections of functional specification of Pathways predicted from KASS automated server. Plot coordinates are differentiated in three layers. First layer specifies Orthology menu of the pathways from prediction profile. Second layer specifies compositional head of pathway ID and third layer is to specific functions of each pathways that corresponds from orthologs.

KEGG analysis showed that most of the expanded genes were involved in signal transduction, protein metabolism, ribosomes, DNA repair mechanisms, and mitochondrial biogenesis, clearly showing that the ricebean genomes represent a huge repository of resources for domesticating crops according to the functional classification of the ricebean genomes. The expanded orthogroups were involved in phenylpropanoid, terpenoid, carotenoid, tropane, piperidine, flavonoid, folate, zeatin, isoquinoline, cutin, sesquiterpenoid, stilbenoid, gingerol, prodigiosin, monoterpenoid, phenazine, novobiocin, isoflavanoid, indole, acridone, benzoxazinoid, clavulanic acid, puromycin, and staurosporine biosynthesis. We observed that 515 genes from contracted gene families clustered in 54 KEGG pathways, including starch, sucrose, purine, pyruvate, inositol, glutathione, glyoxylate, and pyrimidine metabolism (Supplemental Tables 4). The patterns of the genes encoding enzymes distributed in the metabolite pathways indicated the enrichment of hierarchies associated with flavonoid biosynthesis.

#### Transcription factors

Most of the essential domestication genes of crop plants have been shown to encode transcription factors (Borrill *et al.,* 2019). Plants regulate intrinsic gene expression by transcription factors (TFs), transcriptional regulators (TRs), chromatin regulators (CRs) and the basal transcription machinery (Dai *et al.,* 2013). Recruitment of TF-binding domains from transposases or integrases is a recurrent theme in evolution (Chen *et al.,* 2016). A Hidden Markov Model (HMM) profile search revealed many functional factors that influence growth, membrane transport and so forth. In total, 11202 TFs were identified from genes mapped from the assembled reads. In the clusters of gene families, searches were performed for the top clades with the reference genomes.

In total, 15 gene clusters of transcription factors with extensive functions were identified from the total genes predicted from the assembled reads. Comparisons of the total predicted ricebean genes aligned with the mungbean, adzukibean and cowpea genomes (Table 13). RNA1-encoded 60-kDa nucleotide-binding proteins are usually involved in the hybrid systems of plant proteins, and 38% of the total functional homologue count of these proteins was identified in comparison with Vang, 33% for Vrad and 28% for Vung (Carette *et al.,* 2002). For proteins the hexamer motif 5’-ACGTCA-3 (transcription factor HBP-1a), which promotes histone gene binding (Cao *et al.,* 2019), the corresponding values were 39% for ricebean mapped adzukibean [RmA] genes, 33% for ricebean mapped mungbean [RmM] genes and 27% for ricebean mapped cowpea [RmC] genes. For Nitrilase Family Member 2 (NIT2), which cleaves carbon-nitrogen bonds (Urbancsok, Bones and Kissen, 2018), the values were *36% from* RmA*, 33% from* RmM *and 29% from* RmC. For of Opaque endosperm 2 (Opaque-2), which regulates the expression of many members of the zein multigene family of storage proteins (Krishna, Sokka Reddy and Satyanarayana, 2017), the values were 40% for RmA, 34% for RmM and 24% for RmC. 36% for RmA, 33% for RmM and 29% for RmC corresponding to (SEF4) nuclear factor (Zhao *et al.,* 2015) and 35% from RmA, 32% RmM and 31% RmC for common plant regulatory factor-promoting genes (CPRF1), which are regulated by diverse stimuli such as light induction or hormone control (Rügner *et al.,* 2001). The average values were 30% for RmA, 35% for RmM and 33% for RmC for rarer TFs found with variable counts of >1%, including a TEA/ATTS transcription factor [abaA] (Hwang, Chambon and Davidson, 1993), putative chromosomal passenger protein (CPC1) (Carmena *et al.,* 2012), G-box-binding factor (GBF) (de Vetten and Ferl, 1995), facilitative glucose transporter (GT-1), facilitative glucose transporter GT2 (GT-2) (George MT *et al.,* 2015), Sequence-specific single-strand DNA-binding protein-2 (ssDBP-2) (Takase, Minami and Iwabuchi, 1991), TATA-box binding protein-associated factor 1 (TAF-1) (Friedrich *et al.,* 2005), and putative homeodomain-like transcription factor superfamily protein (HOX1a) (Comelli, König and Werr, 1999). Expansion and contraction of TF gene families might influence the regulation of biological functions and trait differences in ricebean (Joshi *et al.,* 2016). The fraction of TF gene families predicted and aligned to the reference genomes indicated an average reduction in genetic distance of 36% in RmA 33% in RmM and 30% in RmC, which was much less than that in ricebean but comparable with that in other legumes.

**Table 13:** Genes involved in transcription factors in rice bean genes mapped from adzuki bean, mung bean and cowpea.

| Gene name expansion | TF | Vang | Vrad | Vung |
| --- | --- | --- | --- | --- |
| 60K protein | 60K protein | 69389 | 60727 | 50914 |
| TEA/ATTS transcription factor | abaA | 2338 | 2366 | 2277 |
| putative chromosomal passenger protein | CPC1 | 1980 | 13824 | 10692 |
| common plant regulatory factor 1-like | CPRF-1 | 15597 | 1830 | 2090 |
| G-box-binding factor | GBF | 159 | 147 | 292 |
| GT1 facilitative glucose transporter GT1 | GT-1 | 1042 | 1010 | 950 |
| GT2 facilitative glucose transporter GT2 | GT-2 | 441 | 369 | 405 |
| hexamer motif 5'-ACGTCA-3 | HBP-1a(c14) | 293575 | 249399 | 204585 |
| Nitrilase Family Member 2 | NIT2 | 49403 | 45274 | 39537 |
| opaque endosperm 2 | Opaque-2 | 25952 | 22118 | 15832 |
| Nuclear factor | SEF4 | 15594 | 14424 | 12870 |
| Sequence-specific single-strand DNA-binding proteins-2 | ssDBP-2 | 352 | 451 | 319 |
| TATA-box binding protein associated factor 1 | TAF-1 | 892 | 930 | 1002 |
| putative homeodomain-like transcription factor superfamily protein | Zmhox1a | 539 | 524 | 369 |

### Genome editing tools in *V. umbellata assembly*

Editing tools have the potential to precisely change the architecture of a genome with the desired precision. These tools can modify the genomic architecture precisely to achieve the required accuracy (Zhang *et al.,* 2018). These tools have been efficiently used for trait discovery and for the generation of crops with high crop yields (Wang *et al.,* 2019). This analysis enables the investigation of evolutionary elements and the molecular functional impact of domestication and breeding in *V. umbellata*. Three important genetically close traits must be addressed during the domestication process make ricebean a viable crop: palatability, latency and an increased rate of flowering (Pattanayak *et al.,* 2019). Nine major functional clusters were derived from 31276 predicted genes with several loci (Dong *et al.,* 2004); i.e., genes that exhibit plasmid specificity, hybridization loci, localization sequences, ORI regions, promoter sequences, reporters, Tags, and terminator sequences.

A total of 33 genes - 7049 loci were identified for vector specificity, whereas the SV40 late 19S internal ribosome showed an abundant distribution (12%) among the total predictions (Table 14). Proportions of VDE, T7 gene10, casp-3, HOI, TEV, RBS Kozak, SV40 int, PHO1, T7 transl RBS, and delta U3 between 4%-6% were identified with an average of 3000 loci The proportions of AraI1I2, incA, kemptide targ, FRT, EK, myr, loxH, loxH; loxP, loxP, and CAP BS ranged from 2-3 %. Proportions of SIN, thromb targ, loxP, loxH, 3 AcPH, Tn7_att, 3xSp1, cosN, LITR, RITR, MODC, Encap, and LAAV-2 ITR of less 1% were prevalent in this study.

Nine main localization genes were identified, among which the MAPKK_NES gene presented the highest frequency in the gene set (i.e., 36% and 3867 loci) (Table 15). Second, an NLS_tandem_repeat was included among the sequences with the highest frequencies (3294 loci, approximately 33%). Calreticulin_targeting loc exhibited a frequency of 13 % and 1301 loci. The values for c-Ha-Ras farnesylation, peroximal target signal1 loc and a region from targeting loc were in the mid-optimal range, from 2-7%, with 758, 417 and 178 loci, respectively. The frequencies of h_a-tubulin loc (34 loci), hCyto_b_actin loc (362 loci) and rhoB_GTPase loc (17 loci) were less than 1%.

SV40 origin and ds origin are two identified sources of ORI and exhibited 298 and 261 loci, respectively (Table 16). One lacZ reporter locus was found. The HOI gene, encoding a post-transcriptional regulator, presented a single locus (Table 17).

Twenty genome editing responsible promoters were screened from the predicted genes. Based on the counts of the total aligned loci, the *lpp* promoter localized to 6225 different regions, corresponding to 20% of the total promoters predicted (Table 18). P5 and amp promoters were distributed at 5611 and 3231 loci, respectively. Ecoli_CI, lac, ARA, tac, T3, Sp6, NEOKAN, PH, T7, PLtet, trc, EM7, CMV, protA, URA3 and CMV2, with frequencies of less than 10%, were distributed at an average of 935 loci.

A total of 31 regulatory elements were present, distributed at 30939 loci. Translational_enhancer_5_prime_UTR (TE5pUTR), p53 response element (p53_RE), Glucocorticoid response element (GRE), Signal transducer and activator of transcription 3 (STAT3_DBS), Interferon-Stimulated Response Element (ISRE), D-glucuronyl C-epimerase (HSE), arabinose operon 2 (araO2), Gamma interferon activation site (GAS) and ethylene responsive element (ERE) sequences were distributed at 3105, 2940, 2870, 2586, 2331, 2313, 1875, 1781 and 1688 loci, respectively whereas other regulatory elements involved in activation, reception and inhibition were distributed at less than 2% of loci; these sequences included the following: activating protein-1 (AP1), arabinose operon 1 (araO1), nuclear factor of activated T-cells (NFAT RE), transcriptional) ulator Myc-like (cMyc DBE), reverse response element (rrnB ATS), retinoic acid receptor DNA binding (RAR DBD), T710RBSL, Tetracycline resistance protein (tetO), nuclear factor kappa B (NFKB RE), transcription factor E2F (E2F DBS), kinase-like protein (mPKA), RNaseIII processing site (RNAseIII PS), RbRE(RbRE), coatomer protein complex subunit alpha (CopA), coatomer protein complex subunit gamma (copG TR), TB, cyclization recombinase (CRE), transporter associated with antigen processing (tap), E-box-directed gene (EGRE), Lactose Operon (lacO) and Tetracycline Inducible Expression (TRE) (Table 19).

Eleven terminators were found at 21996 loci. A total of 5903 loci of synthetic polyadenylation signal terminators, 4862 loci of forward terminators, 4216 loci of transcriptional terminators, 1953 loci of rrnB T2-reverse response typical factor-independent terminator-like sequences, 1292 loci of T7 terminators, 1276 loci of rrnB T1, 924 loci of stem-loop early T7, 9 loci of polyA site-dependent terminators, and 8 loci of yADH1 terminators were identified (Table 20).

#### Genome map distributions

Gene mapping was performed for 414 mb of the assembly, and 31276 genes that were conserved in diverged plant species were found in the nt database. [**a**] A total of 49 mb of genes predicted from the alignment of adzukibean (455 mb), mungbean (459 mb), and cowpea (607 mb); [**b**]105 mb of genes mapped from adzukibean genome; [**c**] 92 mb from mungbean genome and [**d**] 81 mb from mapping cowpea genome. Furthermore, to achieve consistency of the genes deposited in the reference genomes, 49 mb of predicted genes were aligned to the complete reference of *Vigna angularis*, and 105 mb of genes were found, which was 2X times higher than in the predicted gene profiles. In total, 21006 genes in ricebean were identified from adzukibean genome.

#### Synteny clusters

##### Synteny map between ricebean mapped genes from adzukibean (105 mbp) and predicted genes from BLASTX(49 mbp)

Extensive synteny among mapped and assembled reads allowed the integration of their genetic and physical maps. These observations support the hypothesis that detectable synteny exists between the maps and assembly related to divergence, although the conservation is more extensive than in the predicted profiles. The detectable synteny between these 105 mbp and 49 mbp of sequences provides clues for the identification of orthologous blocks associated with functional gene features with increased consistency. At a higher stringency, 80-90% of gene queries with seed weights were found among 653 identified homologs with an assembly that supported the quantification of the complete genome profile of *Vigna umbellata* (Fig 20).

**Fig 20:**
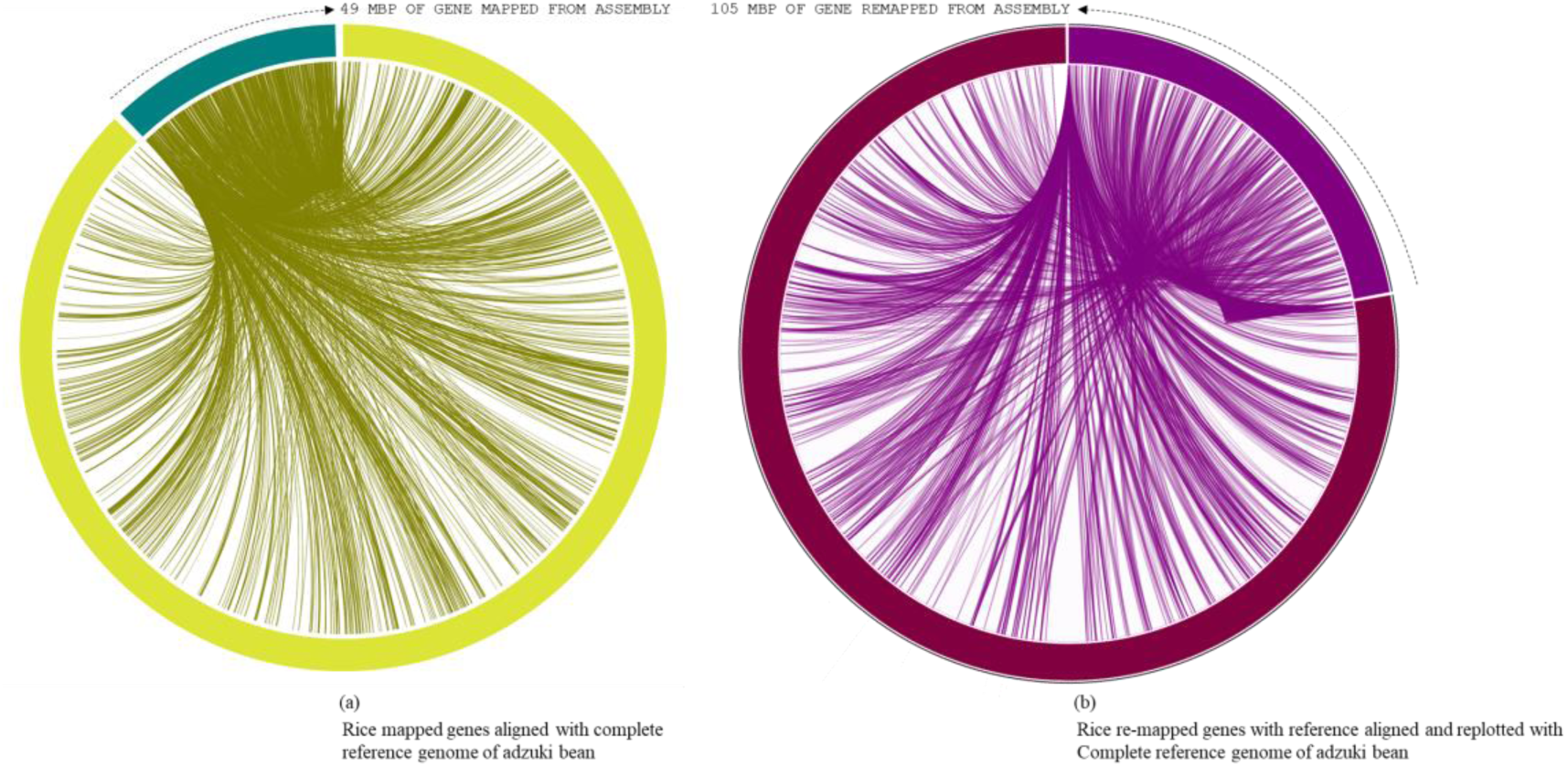
Mapping of predicted genes and genes predicted from remapping with reference genome of Vigna angularis. (a) Gene distribution of 49 mbp of genes with ref genome. (b) Gene distribution of 105 mbp of genes with ref genome.

##### Synteny map between predicted genes from reference genomes

Colinear blocks were calculated, and the flanking regions were plotted using Circa. The homology blocks between RmA (105 mbp), RmM (92 mbp) and RmC (81 mbp) were closer, and high-degree comparisons identified candidates for linkage and highly conserved profiles. To The comparison of total genome positions revealed coverage of >88% of the physical distance of the adzuki bean genome, with 42 conserved synteny blocks, 72% of the mung bean genome and 62% of the cowpea genome, and many synteny microbreak points were found in all three selected reference species. Based on corresponding loci and linear blocks between the genomes, a large number of syntenic blocks between ricebean and adzukibean than ricebean *–* munbean and ricbean *–* cowpea pairs (Fig 21). *Synteny map between predicted genes and mapping against Vigna angularis:* The consistency of the genes and conserved profiles were extensively studied by plotting syntenic data from Mauve alignment. In total, among 24516 genes from *Vigna angularis*, 78% of the genes were identified from the BLASTx gene prediction, and 87% were identified from remapping with adzukibean. These values were calculated based on genetically anchored colinear blocks. The comparison of remapped ricebean genes versus single-copy genes in adzukibean showed improved visualization of synteny with single-copy genes (Fig 22). *Synteny map between ricebean mapped genes and complete chromosome sets of Vigna angularis:* The homolog richness of the *Vigna umbellata* genes mapped against 11 chromosomes of *Vigna angularis* was evaluated (Fig 23). The alignment results revealed that Chr1, Chr3 and Chr9 were the chromosomes showing the high functional chromosomal alignment, for which the contributed genome size was greater than 40 mbp. The conservation of synteny between Chr7, Chr2, Chr4 and Chr10 was closer to >30 mbp, and Chr8, Chr5 and Chr11 constituted a group with low syntenic (micro) matches. The lowest conservation of homologs was found in chromosome 6, which corresponded to less than 19 mbp of the total genome size. The closer comparison of the high conservation of macro- to microsynteny between mapped genes and *Vigna angularis* revealed a difference of by only approximately 10 mbp of syntenic localisation. The occurrence of prominent duplication events has been speculated because of the extent of collinearity blocks, which are more conserved and functional in relation to *Vigna umbellata*.

**Fig 23:**
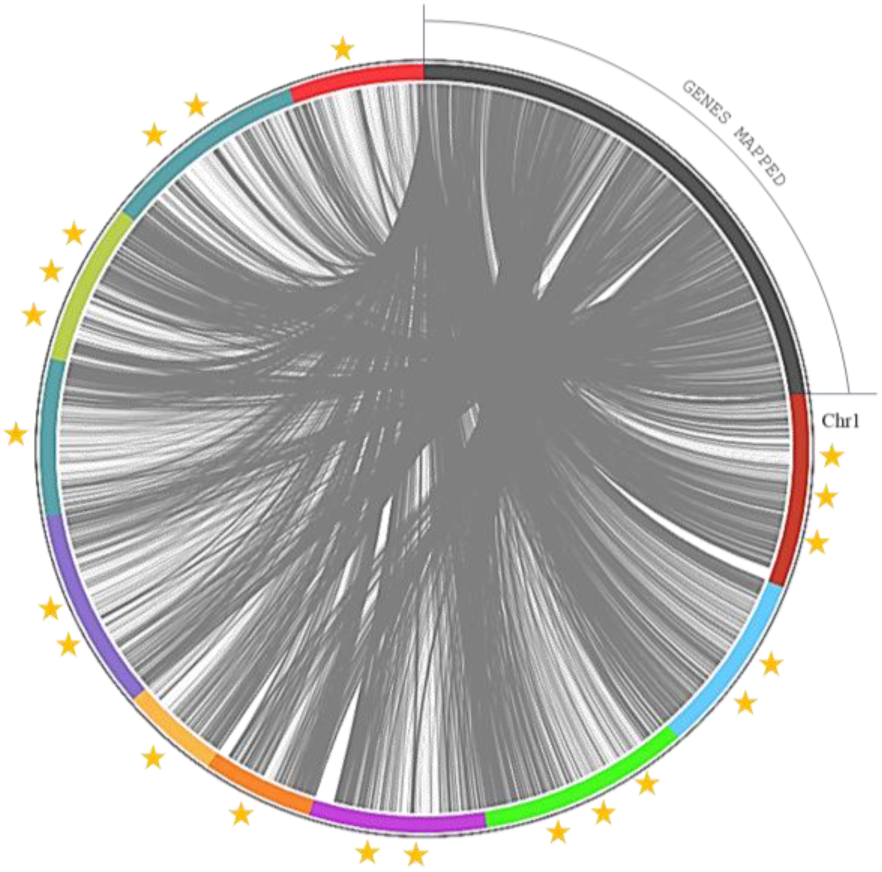
Mapped genes rice bean are aligned with angularis (genetically closely related species). Three star rated chromosome shares 40 mbp of genome size. Two star rated shares >30 mbp os genome size and single star rated chromosomes are 19 mbp of genome map.

### Comparison of 32 Leguminous plant Genomes with ricebean genomes

More than 14000 species are reported as leguminous crops, including both edible and non-edible species (Ng, 2012). Leguminous annuals, perennials, shrubs, vines and trees have adapted to a range of growing conditions around the world (Latef and Ahmad, 2015). Thirty-two species are edible and are categorized as critically important economic food crops (Graham and Vance, 2003). Derived biological statistics from the FAO were used to list crops according to their economic significance based on the production range, postanthesis period and photoperiods of individual species (Pooran *et al.,* 2018). We selected 32 species that are edible, economically important, underutilized crops or staple crops for comparison with the sequenced rice bean genome. The selected grains are intentionally grown for the harvesting of mature seeds, and these species are small fractional leguminous crops with lower production (Clement *et al.,* 2019) than the complete group, such as *Psophocarpus tetragonolobus*, *Lupinus albus*, *Vigna unguiculata*, *Vigna mungo*, *Vigna aconitifolia*, *Vigna radiata*, *Lupinus angustifolius*, *Lupinus luteus*, *Mucuna pruriens*, *Vigna subterranea*, *Cyamopsis tetragonoloba*, *Cajanus cajan*, *Pachyrhizus erosus*, *Lens culinaris*, *Lupinus mutabilis*, *Lablab purpureus*, *Vicia faba*, *Vigna angularis*, *Phaseolus coccineus*, *Vicia sativa*, *Cicer arietinum*, *Canavalia gladiata*, *Dolichos lablab*, *Phaseolus lunatus*, *Lupinus perennis*, *Phaseolus acutifolius*, *Pisum sativum*, *Glycine max*, *Arachis hypogaea*, *Phaseolus vulgaris* and *Canavalia ensiformis*. Among the 31 species selected, the complete genomes have been reported for 10 species (6 chromosomal annotations + 4 raw annotations) (Table 26), and for the other 21 species 44% of ribosomal genes and other genes with various functions have been reported (Table 24).

**Table 24:** Annotation percentage of Leguminous plants prevailing complete genome profile from NCBI Genome.

| Species | No. of sequence | No. of seq aligned | % of seq. aligned |
| --- | --- | --- | --- |
| *Vigna angularis* | 72 | 35 | 48.61 |
| *Phaseolus vulgaris* | 1620 | 697 | 43.02 |
| *Vigna radiata* | 79 | 27 | 34.18 |
| *Phaseolus coccineus* | 6 | 2 | 33.33 |
| *Glycine max* | 696 | 213 | 30.60 |
| *Cajanus cajan* | 997 | 143 | 14.34 |
| *Vicia faba* | 181 | 25 | 13.81 |
| *Vigna unguiculata* | 80 | 7 | 8.75 |
| *Arachis hypogaea* | 43 | 1 | 2.33 |
| *Lupinus angustifolius* | 83 | 1 | 1.20 |
| *Cicer arietinum* | 98 | 1 | 1.02 |
| *Phaseolus acutifolius* | 1 | 0 | 0.00 |
| *Pisum sativum* | 2 | 0 | 0.00 |
| *Vigna aconitifolia* | 2 | 0 | 0.00 |

**Table 26:** List of genome composition and distribution of selected leguminous plants for rice bean genome alignment.

| Species names | Common names | Median total length | Median GC count | No. of chromosomes | Size in bytes |
| --- | --- | --- | --- | --- | --- |
| *Arachis hypogaea* | Nuts | 2551.68 | 36.3706 | 20+2 | 2.5 GB |
| *Cajanus cajan* | Peas | 620.626 | 33.6052 | 11+2 | 620 MB |
| *Cicer arietinum* | Peas | 510.877 | 32.4428 | 8+2 | 510 MB |
| *Glycine max* | Bean | 997.663 | 35.1918 | 20+3 | 997 MB |
| *Lupinus angustifolius* | Lupins | 557.909 | 33.5664 | 20+1 | 557 MB |
| *Mucuna pruriens* | Pulses | 397.042 | 31.4 | 1WGS | 397 MB |
| *Phaseolus coccineus* | Dry beans | 371.086 | 32.1 | 1WGS | 371 MB |
| *Phaseolus vulgaris* | Dry beans | 535.413 | 36.8761 | 11+1 | 535 MB |
| *Pisum sativum* | Dry peas | 4257.93 | 37.7 | 1WGS | 4.2 GB |
| *Vicia faba* | Dry broad beans | 80.3627 | 38.9 | 1WGS | 80 MB |

The complete genome (Fig 27) aligned with *Arachis hypogaea* genome (2.5 GB genome size), *Cajanus cajan* genome (620 MB), *Cicer arietinum* (510 MB), *Glycine max* genome (997 MB), *Lupinus angustifolius* genome (557 MB), *Mucuna pruriens* genome (397 MB), *Phaseolus coccineus* genome (371 MB), *Phaseolus vulgaris* genome (535 MB), *Pisum sativum* genome (4.2 GB) and *Vicia faba* genome (80 MB), which included the total chromosomal data and assembly annotations, to understand the genetic makeup and versatility of the genome structure on the basis of the genes predicted from ricebean assembly (Table 28).

**Fig 27:**
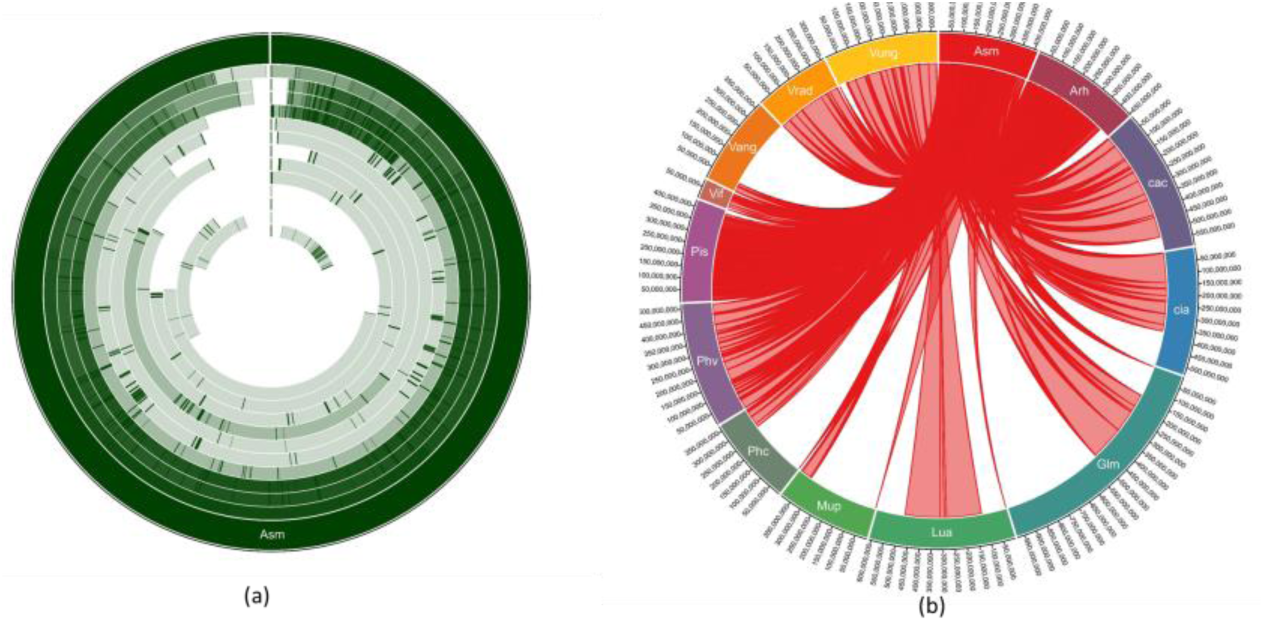
Circular distributions rice bean genome mapping with Arachis hypogaea, Cajanus cajan, Cicer arietinum, Glycine max, Lupinus angustifolius, Mucuna pruriens, Phaseolus coccineus, Phaseolus vulgaris, Pisum sativum and Vicia faba. (a) circular distribution and (b) Connective distributions of rice bean genome assembly.

**Table 28:**
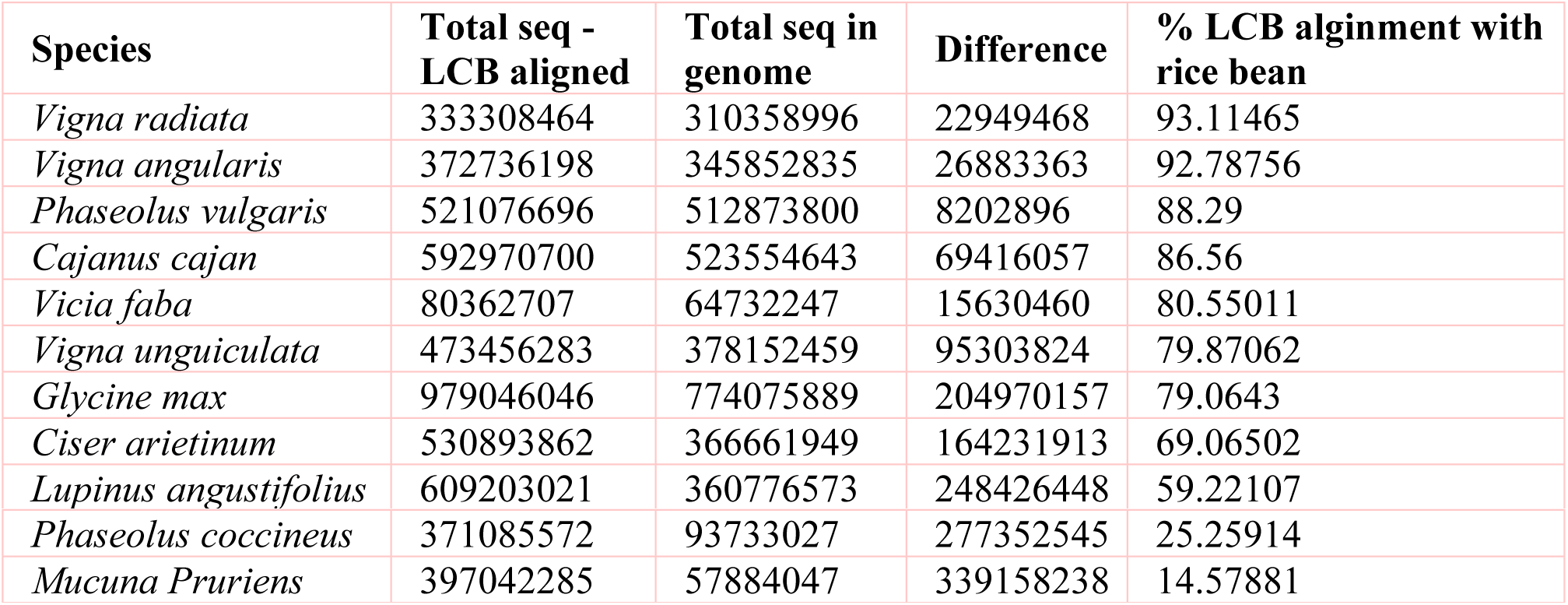
Percentage of complete genome alignment from all selected species having full genome annotation profile.

These crops were selected based on their photoperiod and postanthesis period to improve multi-parent advance generation intercross populations (MAGIC) for studying the genetic structure of rice bean traits and improving the breeding of rice bean accessions (Bandillo *et al.,* 2013). The production range of these crops is influenced by and fundamentally dependant by their flowering potential and photoperiod sensitivity. Rice bean requires a photoperiod with 12 hrs of daylight to flower between 120-160 days in a year (Joshi K.D *et al.,*2007). The photoperiod range comprises much overlap compared with other species, and the species selected for this study is predicted to require a longer photoperiod than other crops but to present an equal nutritional potential. In the production range of rice bean with a prevalent 12 hr photoperiod, 30000 tons of production can be achieved in a year (Dhiman *et al.,* 2010). According to Singh *et al.,* rice bean is a crop with a smaller production range than other species requiring a 12 hr photoperiod, such as *Vigna unguiculata*, *Mucuna pruriens*, *Vigna subterranea*, *Lens culinaris*, *Lupinus mutabilis*, *Vigna angularis*, *Phaseolus coccineus*, *Canavalia gladiata*, *Lupinus perennis*, *Phaseolus acutifolius* and *Pisum sativum*. The production of these species ranges from 430000 tons to 2800000 tons. The postanthesis profile ranges from 70 days to 150 days of the flowering period (Table 25).

**Table 25:**
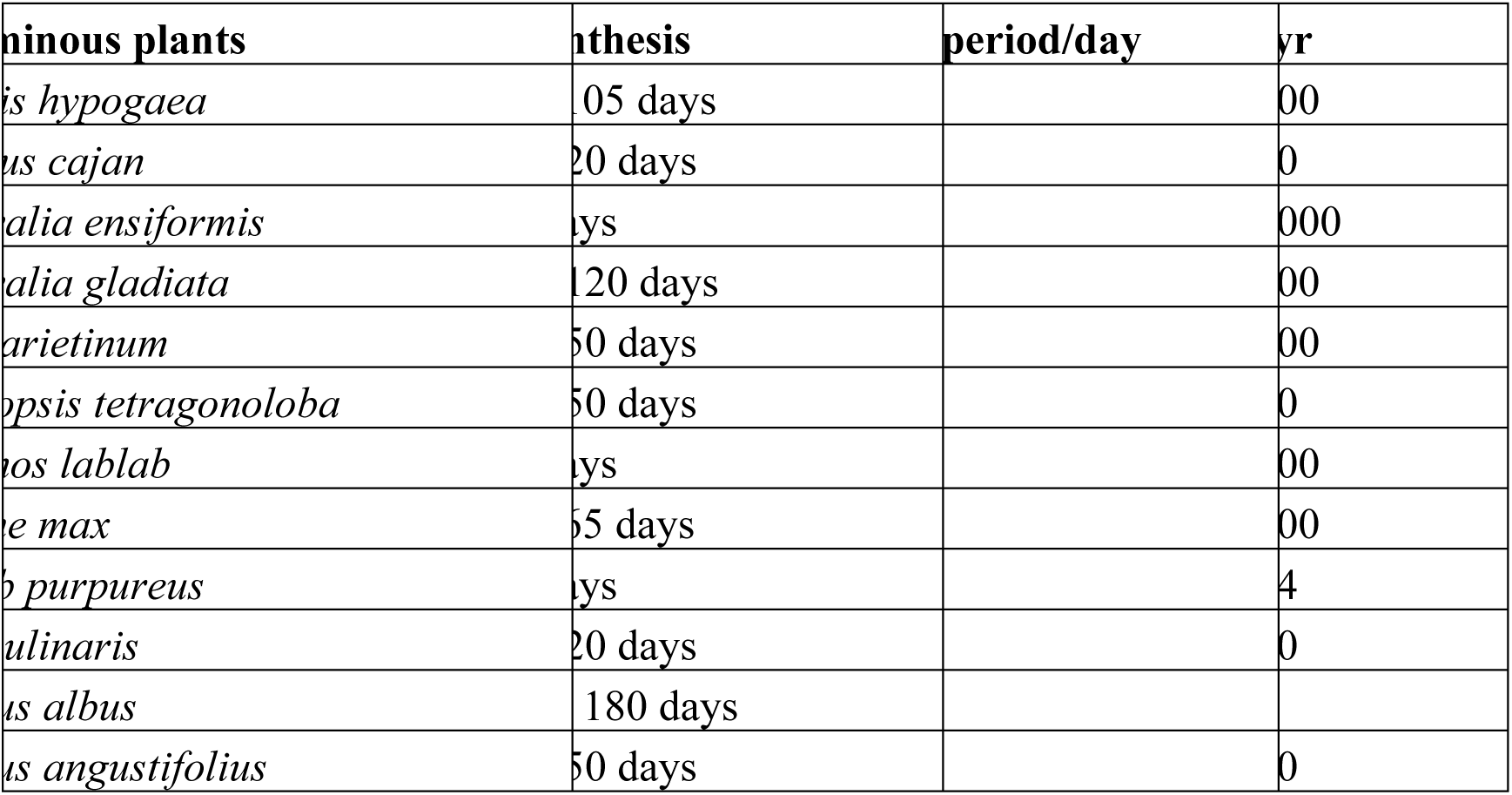
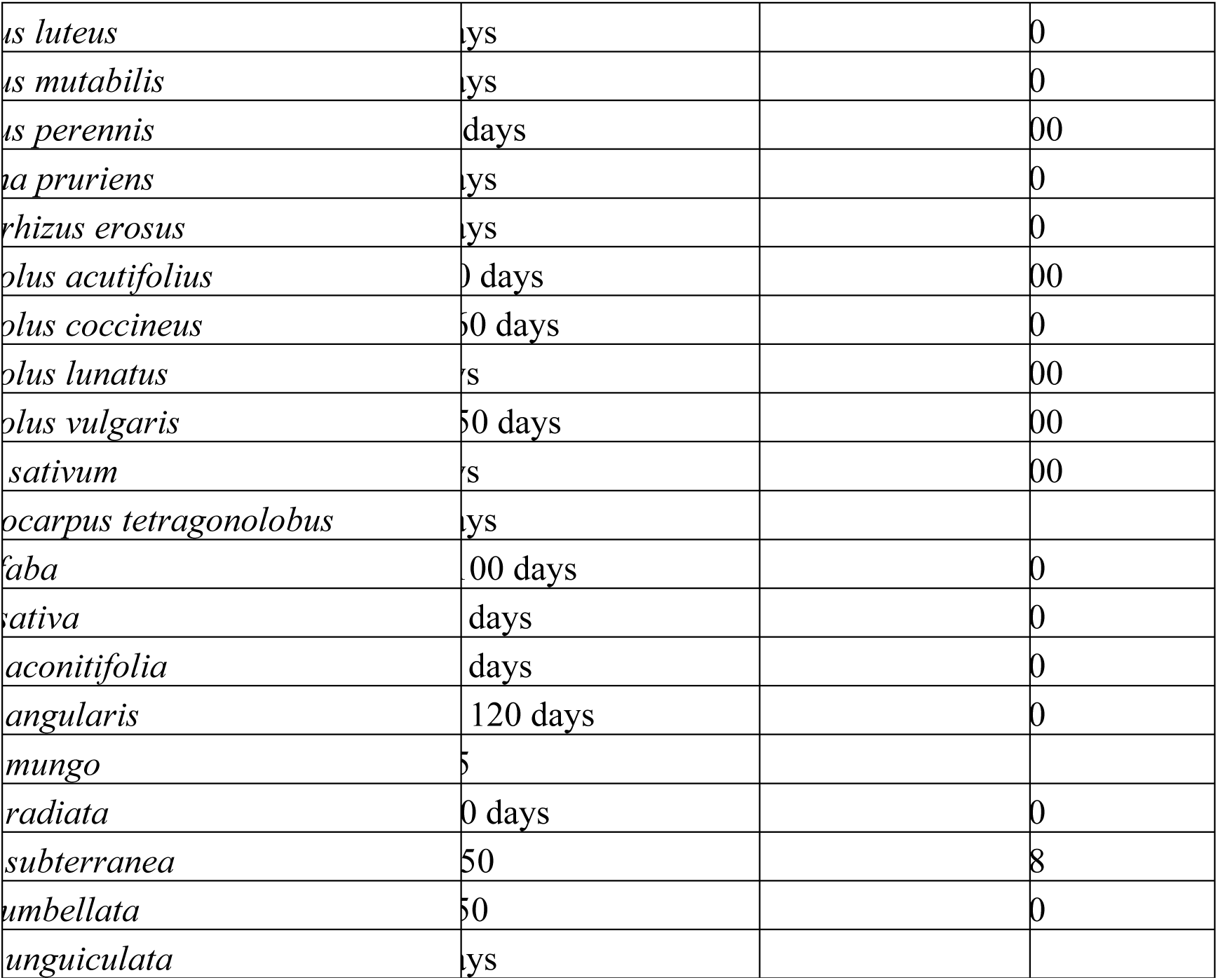
Distributions of Postanjthesis, photoperiod per day and productive range per year for all selected leguminous plants.

Phylogenetic evidence based on ribosomal genes reported from selected species considering the rice bean data as an outgroup was used for the comparison of the genetic distances. (a) The scale range was calculated as 0.7 based on 55 nodes and 28 edge nodes. A distance of 0.01 was calculated for rice bean in the outgroup range in which *Vigna angularis* shared the closest monophyletic distance (0.02) (Fig 26); (b) *Vigna angularis* and *Lupinus angustifolius* shared the same nodes located close to rice bean. The total length of the ribosomal RNA genes varied from 50 bp – 133 bp.

**Fig 26:**
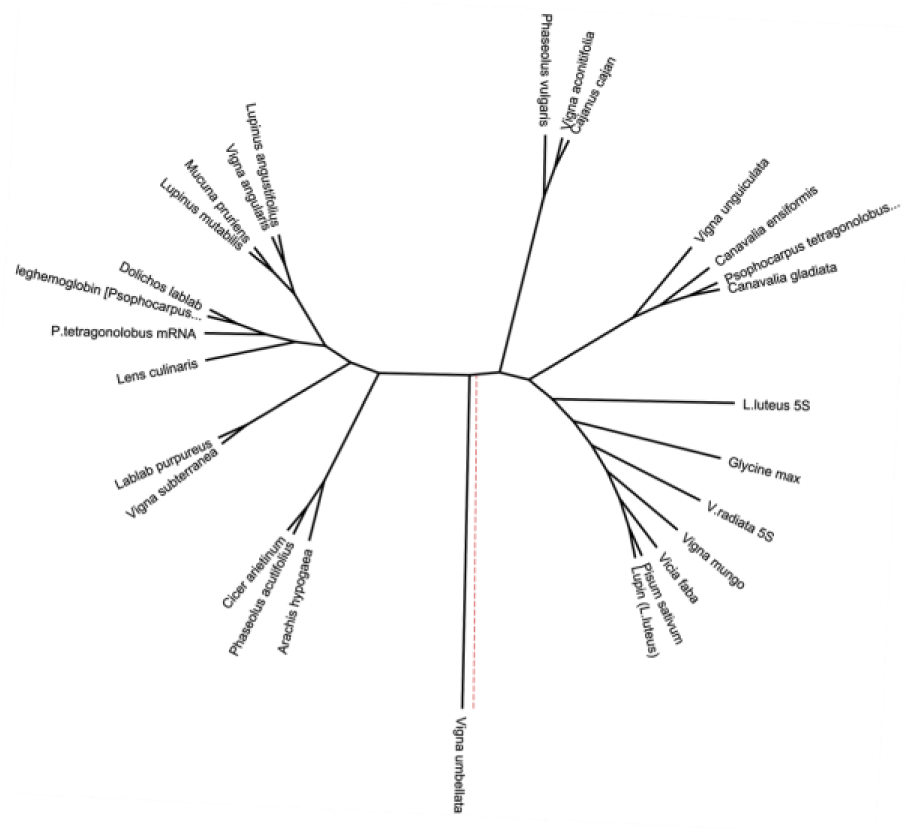
Phylogenetic construction of selected 32 species of leguminous plant genomes outgrouped by Vigna umbellata.

Complete CDS were organized for alignment with the assembled rice bean sequence. Thirteen species, including *Vigna angularis*, *Vigna radiata* and *Vigna unguiculata*, *Mucuna pruriens*, *Phaseolus coccineus*, *Vicia faba* and *Pisum sativum*, lacked a CDS annotation report. In total, 9 complete CDS sequences were aligned based on the LCB count, and the top 5 alignments shared more than 99% CDS alignment. The *Vigna angularis* CDS length covered approximately 99.8% of the total length, followed by that of *Vigna radiata* at 99.7%, *Vigna unguiculata* at 99.2%, *Phaseolus vulgaris* at 99.27% and *Cajanus* at 99.2%, which were all aligned over more than 99% of the profile. *Ciser arietinum*, *Lupinus angustifolius*, *Glycine max* and *Arachis hypogea* showed between 70% - 95% CDS alignment. The *Vigna angularis* total LCB bp count was bp 44,382,592 out of 44,563,217 bp; the *Vigna radiata* total LCB bp count was 50,174,463 out of 50,285,496 bp; the *Vigna unguiculata* total LCB bp 58,335,559 count was out of 58,429,249 bp; the *Phaseolus vulgaris* total LCB bp 40,764,835 count was out of 41,064,196 bp; and the *Cajanus cajan* total LCB bp 31,429,340 count was out of 31,679,668 bp. The other four species, *Ciser arietinum*, *Lupinus angustifolius*, *Glycine max* and *Arachis hypogea*, shared 41,644,172 out of 43,814,935 bp, 64,299,632 out of 69,566,077 bp, 87,887,691 out of 98,033,242 bp and 137,827,886 out of 194,742,024 bp respectively (Fig 28).

**Fig 28:**
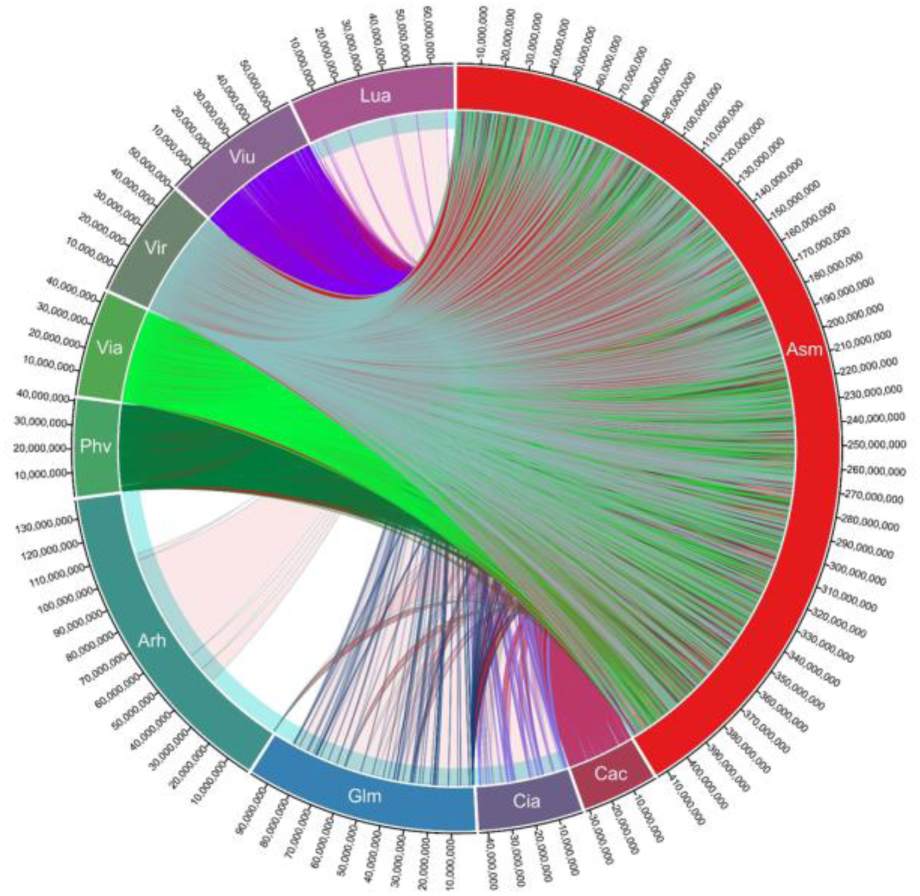
Complete CDS coverage map of Arachis hypogaea, Cajanus cajan, Cicer arietinum, Glycine max, Lupinus angustifolius, Mucuna pruriens, Phaseolus coccineus, Phaseolus vulgaris, Pisum sativum and Vicia faba with rice bean assembled sequence.

The genomic data for the selected leguminous plants that for which there were not available complete genome annotations were downloaded (until July 2019 from NCBI FTP) and aligned with genes predicted from the rice bean assembly. The total number of sequences deposited from each species were subjected to BLAST searches against genes predicted from rice bean, and the total number of sequences that aligned over more than 300 bp was determined. A total of 61.54% (39 seq out-of 24 total seq) of the total *Pachyrhizus erosus* sequences were aligned with rice bean genes, and the corresponding values were 43.71% (789 seq out of 1805 total seq) for *Dolichos lablab*, 43.71% (789 seq out of 1805 total seq) for *Lablab purpureus*, 37.06% (391 seq out of 1055 total seq) for *Vigna mungo*, 34.13% (242 seq out of 709 total seq) for *Lupinus luteus*, 29.36% (315 seq out of 1073 total seq) for *Vigna aconitifolia*, 26.64% (7017 seq out of 26343 total seq) for *Lens culinaris*, 19.34% (1908 seq out of 9864 total seq) for *Lupinus albus*, 19.25% (3190 seq out of 16569 total seq) for *Cyamopsis tetragonoloba*, 19.23% (10 seq out of 52 total seq) for *Canavalia ensiformis*, 14.67% (11 seq out of 75 total seq) for *Psophocarpus tetragonolobus*, 9.68% (3 seq out of 31 total seq) for *Vigna subterranea*n, 8.32% (82 seq out of 986 total seq) for *Phaseolus lunatus*, 7.89% (3 seq out of 38 total seq) for *Canavalia gladiate*, 5.89% (67 seq out of 1138 total seq) for *Vicia sativa*, 5.88% (4 seq out of 68 total seq) for *Lupinus mutabilis* and no sequence match with *Lupinus perennis* (Table 27).

**Table 27:**
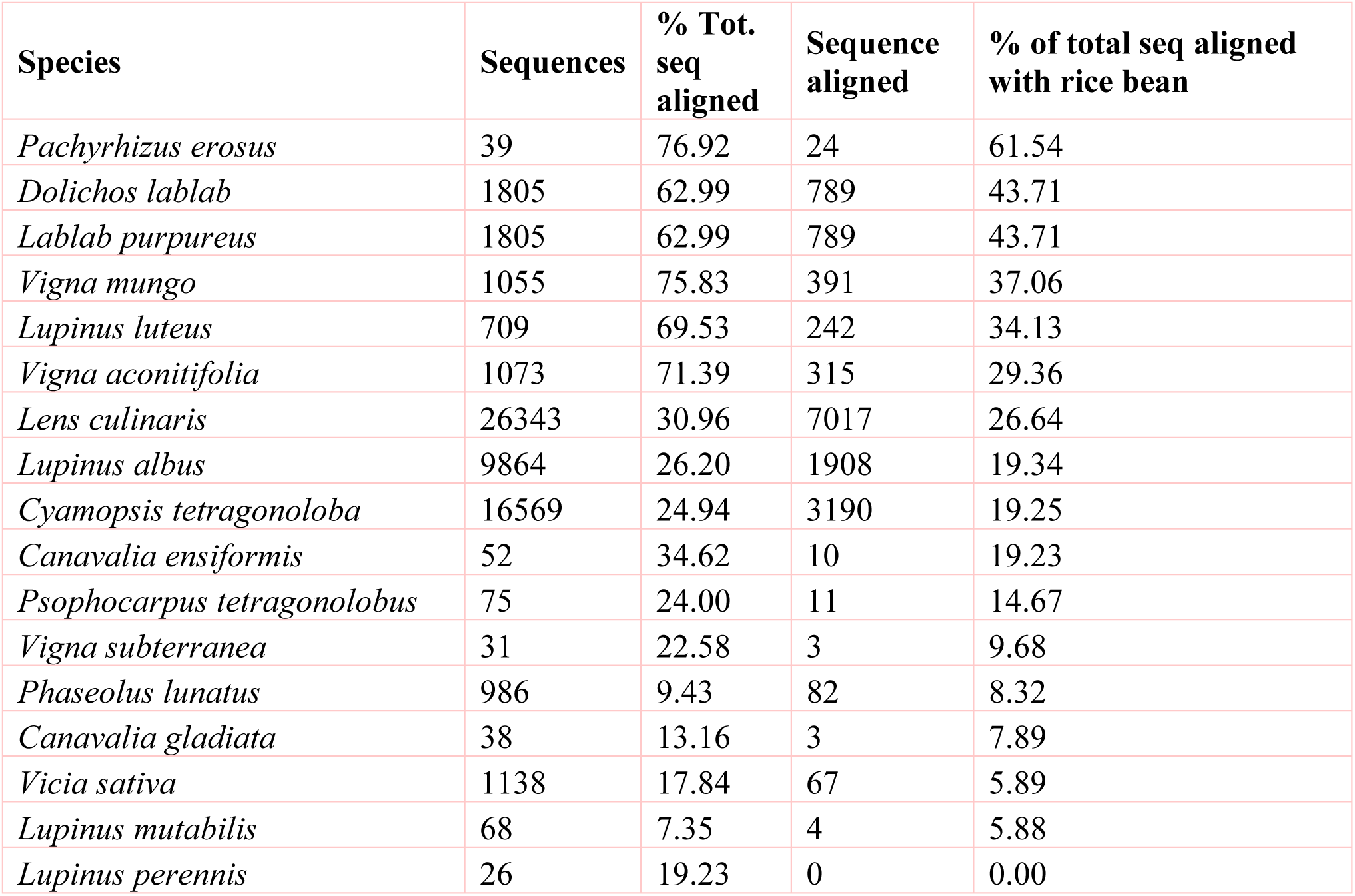
Percentage distribution of rice bean genes aligned with genes retrieved from actual deposition at NCBI Nucleotide database.

| Species | Sequences | % Tot. seq aligned | Sequence aligned | % of total seq aligned with rice bean |
| --- | --- | --- | --- | --- |
| *Pachyrhizus erosus* | 39 | 76.92 | 24 | 61.54 |
| *Dolichos lablab* | 1805 | 62.99 | 789 | 43.71 |
| *Lablab purpureus* | 1805 | 62.99 | 789 | 43.71 |
| *Vigna mungo* | 1055 | 75.83 | 391 | 37.06 |
| *Lupinus luteus* | 709 | 69.53 | 242 | 34.13 |
| *Vigna aconitifolia* | 1073 | 71.39 | 315 | 29.36 |
| *Lens culinaris* | 26343 | 30.96 | 7017 | 26.64 |
| *Lupinus albus* | 9864 | 26.20 | 1908 | 19.34 |
| *Cyamopsis tetragonoloba* | 16569 | 24.94 | 3190 | 19.25 |
| *Canavalia ensiformis* | 52 | 34.62 | 10 | 19.23 |
| *Psophocarpus tetragonolobus* | 75 | 24.00 | 11 | 14.67 |
| *Vigna subterranea* | 31 | 22.58 | 3 | 9.68 |
| *Phaseolus lunatus* | 986 | 9.43 | 82 | 8.32 |
| *Canavalia gladiata* | 38 | 13.16 | 3 | 7.89 |
| *Vicia sativa* | 1138 | 17.84 | 67 | 5.89 |
| *Lupinus mutabilis* | 68 | 7.35 | 4 | 5.88 |
| *Lupinus perennis* | 26 | 19.23 | 0 | 0.00 |

The complete genome sequences of *Phaseolus coccineus*, *Vicia faba*, *Mucuna pruriens*, *Glycine max*, *Lupinus angustifolius*, *Vigna unguiculate*, *Vigna radiata*, *Vigna angularis*, *Ciser arietinum*, *Phaseolus vulgaris* and *Cajanus cajan* were subjected to the calculation of Locally Colinear Bocks with the aligned regions of rice bean assembly. We found that *Vigna angularis*, *Vigna radiata* and *Vigna unguiculata* presented the closest genetic profiles based on the annotation of the assembly with the corresponding genomes. The assembly sequence was aligned with the sequences of other legume plants with reported complete genome sequences. The resultant alignment percentages were 88.29% (total LCB aligned 512 mbp sequence of 521 mbp) for *Phaseolus vulgaris*, 86.56% (523 mbp of 592 mbp) for *Cajanus cajan*, 80.55% (64 mbp of 80 mbp) for *Vicia faba*, 79.06% (74 mbp of 979 mbp) for *Glycine max*, 63% (530 mbp of 366 mbp) for *Cicer arietum*, 53% (360 mbp of 609 mbp) for *Lupinus augustifolius*, 25.25% (93 mbp of 371 mbp) for *Phaseolus coccineus* and 14.57% (57 mbp of 397 mbp) for *Mucuna pruriens* (Fig 27; Table 28).

### Genome alignment of 14 metabolically and pharmaceutically active medicinal plant genomes

Fourteen taxonomically diverse and pharmaceutically important medicinal plant genomes reported by the Medicinal Plant Genomics Resource, Michigan State University, United States, were aligned with the rice bean genome to identify an unparalleled resource for human health. For *Rauvolfia serpentine*, the greatest linear block of the medicinal genome aligned with rice bean corresponded to approximately 43.35% (161 mbp) (Fig 29) of the total rice bean sequence alignment and 89.63% (179 mbp out of 414 mbp) of the total genome size of *Rauvolfia serpentine*. For the other medicinal plant genomes, the alignment percentages are listed in Table 31.

**Fig 29:**
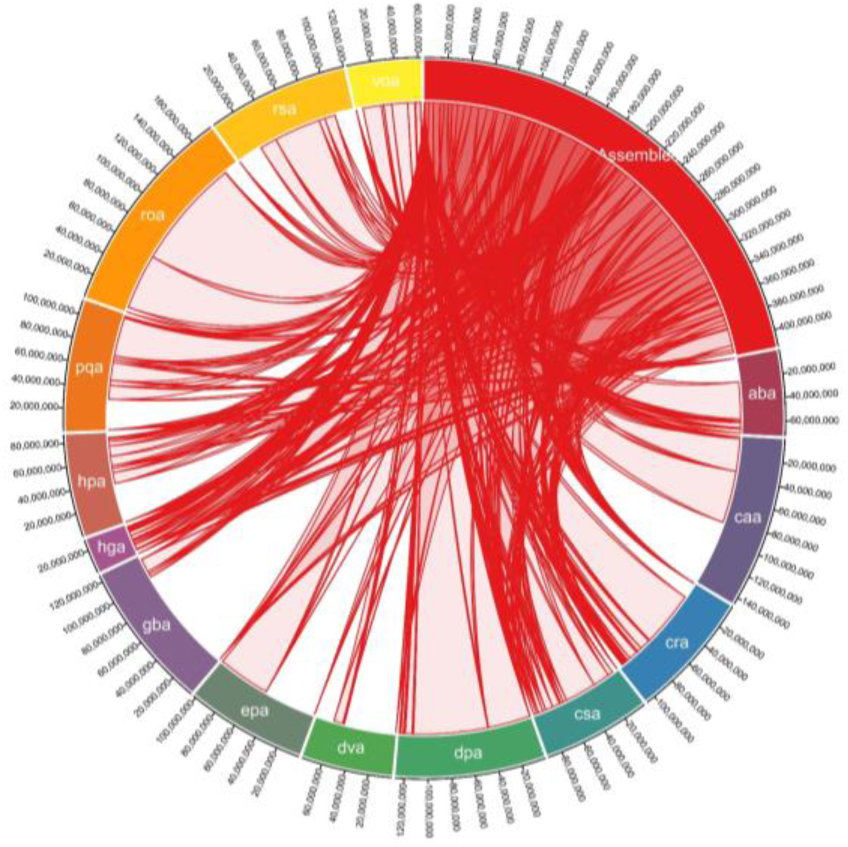
Medicinal genome comparison of Rauvolfia serpentine, Camptotheca acuminate, Digitalis purpurea, Ginkgo biloba, Rosmarinus officinalis, Panax quinquefolius, Cannabis sativa, Echinacea purpurea, Catharanthus roseus, Hypericum perforatum, Dioscorea villosa, Atropa belladonna, Valeriana officinalis and Hoodia gordonii species with rice bean assembled reads.

**Table 31:** List of 14 vital medicinal plants aligned with rice bean complete assembly.

| Species Notation | Species | Tot bp blocks aligned | Total length of the genome | % of tot LCB | % with rice bean genome |
| --- | --- | --- | --- | --- | --- |
| roa | *Rauvolfia serpentina* | 161099773 | 179746725 | 89.625985 | 43.3 |
| caa | *Camptotheca acuminata* | 81953592 | 145899039 | 56.171441 | 35.2 |
| dpa | *Digitalis purpurea* | 126940164 | 127551451 | 99.520753 | 30.8 |
| gba | *Ginkgo biloba* | 19139196 | 125927755 | 15.198553 | 30.4 |
| rsa | *Rosmarinus officinalis* | 77676714 | 120161978 | 64.643338 | 29 |
| pqa | *Panax quinquefolius* | 81232884 | 112315063 | 72.325903 | 27.1 |
| cra | *Cannabis sativa* | 88845168 | 107796557 | 82.4193 | 26 |
| epa | *Echinacea purpurea* | 50867288 | 102468088 | 49.642078 | 24.7 |
| csa | *Catharanthus roseus* | 85738509 | 92280572 | 92.910682 | 22.3 |
| hpa | *Hypericum perforatum* | 44412301 | 87389108 | 50.821323 | 21.1 |
| dva | *Dioscorea villosa* | 10296467 | 77321611 | 13.316416 | 18.6 |
| aba | *Atropa belladonna* | 49133508 | 72725089 | 67.560602 | 17.5 |
| voa | *Valeriana officinalis* | 55758812 | 62865151 | 88.695901 | 15.2 |
| hga | *Hoodia gordonii* | 23869178 | 29731667 | 80.282004 | 7.17 |

### Gene pooling for rice bean domestication and crop improvement

Flowering potential, palatably-indexed genes, stress response, and disease resistance genes were the four gene families that were taken into account for screening and expression analysis in the rice bean genome (Fig 30). *Palatability-specific gene pooling:* A total of 27 genes (ASB2, ASA2, PRTA, PAI2, COMT, CHS, CHS-17, CHI, BAN, UFGT, LDOX, 5MAT, HIDH, IFS2, FLS1, IFR, IFMT, IFH, IFGT, BCKDHA, ILMT, REM, ATB, ROMT, SPDSYN2, SDT, and SHT) corresponding to closer nodes for bitter metabolite functions (Dagan-Wiener *et al.,* 2018) that are crucial to palatably-indexed which activates Taste receptors in human gustations, were retrieved and analysed in the selected five ricebean varieties (PRR-2010-2, PRR-2011-2, RBHP-307, RBHP-104, and VRB-3) along with Glycine max as reference transcriptome sample (SRA : SRX6788895). We found 3 loci (scaffold specific) of 27 specific genes expressed in selected reference (*Gmax*). *Flowering specific gene pooling:* 11 flowering potential genes from selected legume crops were subjected to BLAST search against ricebean draft genome from the assembly and analysed for transcript expressions; found 3 genes that are responsible for late flowering, is expressed high in Glycine max and 8 genes are not expressed in ricebean that encounters the commitment of late flowering of ricebean. *Disease resistance gene pooling:* the EDR-like, ADR, TAO, CSA RPP, RPM, and TMV \ gene families were pooled from mung bean, adzuki bean and cowpea. In total, 331 genes were found, and 205 loci were aligned to the rice bean genome. Coding region domains were rich in the alignments of RPM1 isoforms and Enhanced disease resistance 2-like and RLM3 – like isoforms, whose lengths from the rice bean scaffolds were distributed between 3000-4000 bp. 4% of the RGA1, RGA2 and RGA4 gene families, 15.68% of the RPP13 gene families, 10.2% of the EDR1 and 2 families, 7.8% of the RRP13 and RPM1 isoforms, and other gene families such as ADR2, TAO1, homologs of resistance proteins, RLM3, DSC1, LAZ5, RML1A, RPP8, TMVN, CSA1 and NPR1 accounted for 15% of the disease resistance gene composition in annotated ricebean draft genome. *Stress-responsive genes:* 23 Stress-responsive gene families found in other leguminous plants, were aligned with ricebean predicted genes. 5 gene families out of 23 selected genes [Stress enhanced protein 1 (SEP1-), Universal stress protein PHOS32 precursor (PHOS32), Stress enhanced protein 2 (SEP2-), and Stress responsive alpha-beta barrel domain-containing protein (GSU2970)] were found upregulated in ricebean, wherein it is downregulated in Glycinemax. The list of stress sequence coverage, expressions and the species distribution are provided in Table 29.

**Fig 30:**
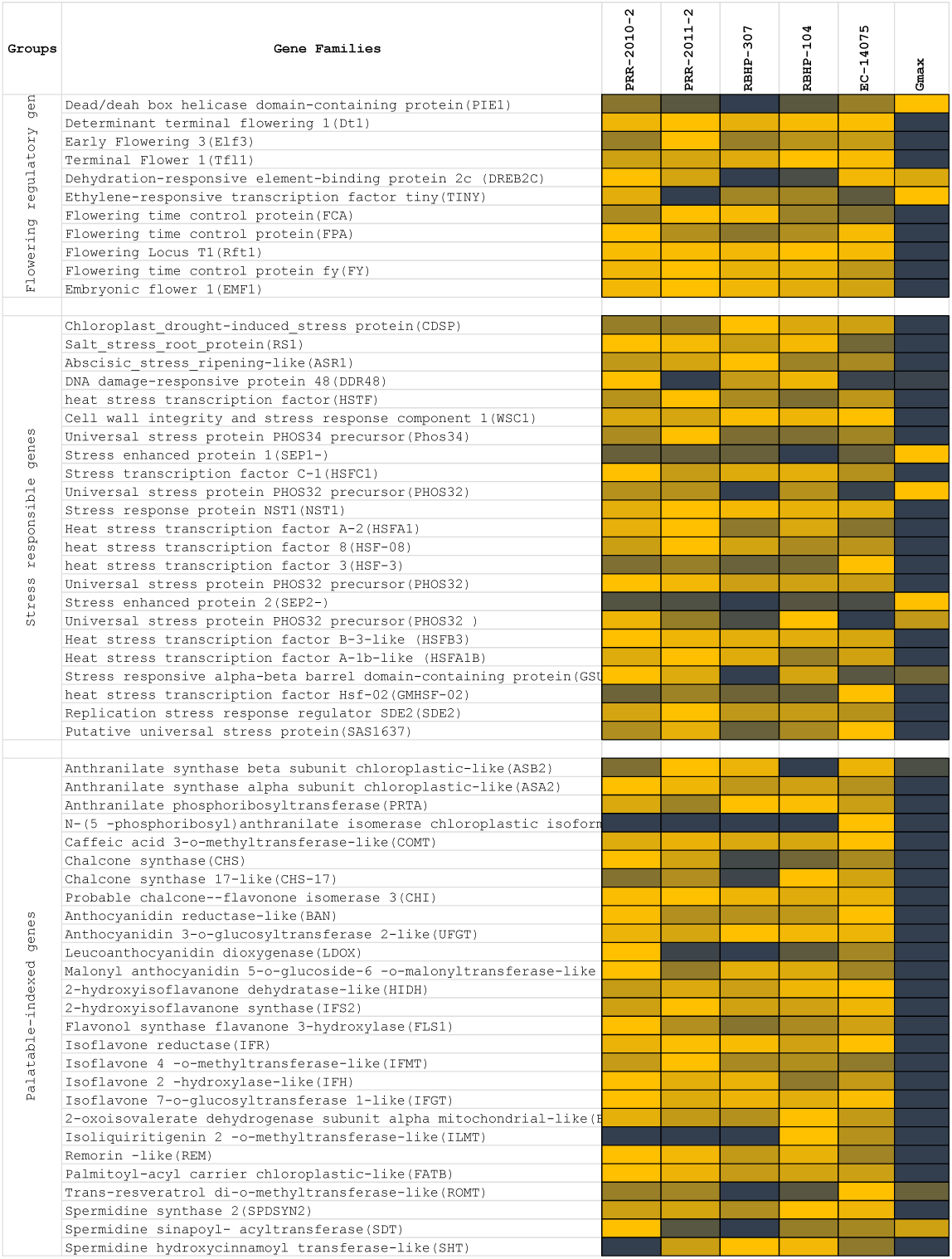
List of genes’ expression selected for late flowering, ligands produced by palatably - indexed genes that activates human bitter taste receptors, stress response genes, and disease resistance genes. RNA expressions of five ricebean varieties (log10 of RPKM of total read counts) and denovo assembled RNA expressions of Glycine max was compared for regulations of each functional genes selected for expression profile analysis.

**Table 29:**
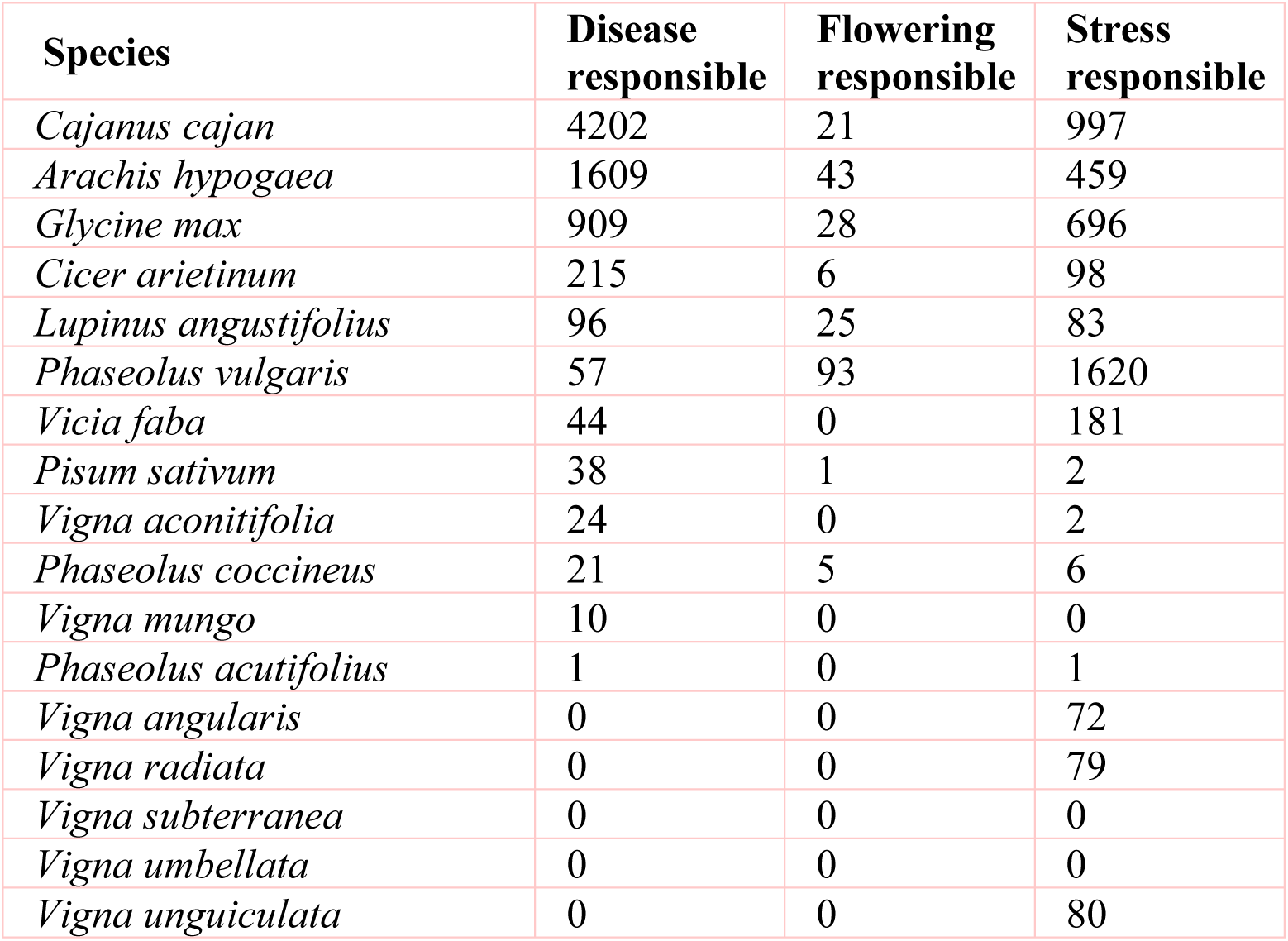
Total no. of genes retrieved from NCBI nucleotide sequence database with respect to Disease, flowering and stress responsible gene families.

## DISCUSSION

Rice bean is less familiar among Asiatic *Vigna* varieties but presents higher potential value than adzuki bean, mung bean and cowpea, which are in the same genetic linkage map in the genome profile. Rice bean has long been considered a food security crop of small and marginal farmers in Southeast Asia. A comparative and functionally curated library was developed from the draft genome sequence of *Vigna umbellata* from multiple compendia. This study showed that the crop species *Vigna umbellata* is relatively rich in biochemical pathways, enzyme distribution and reaction types, active phytocompound-specific hub genes, genome organization, composition of protein-coding genes, the relative composition of non-coding RNAs, etc.

The domestication and subsequent breeding of *Vigna umbellata* focusing on its palatability and flowering potential, stress responsive genes, and disease resistance gene families have had inconsistent effects on its genetic diversity. Most varieties of ricebean are highly photoperiod sensitive and are therefore late flowering show strong vegetative growth when grown in the subtropics. The induced mutation of genes on the basis of the present study could transform a climbing perennial that develops axillary inflorescences during a juvenile stage of many years into a compact plant with rapid terminal flower and fruit development. Expressions of important flowering-related gene families are found significant in five selected varieties of ricebean by positioning read mapping values with *Glycine max*. The Early Flowering 3(Elf3) gene found upregulated in rice bean varieties that would increases functional factors of transcriptional coregulation activity (corepressor complex), because of long-term interaction with Phytochrome B and nematode feeding. Determinante Stem 1 (DT1) negatively regulates and expressed high in the ricebean and downregulated in controlgroup shows delay period of flower development from the vegetative to reproductive phase transition in the meristem. PIE (Photoperiod-Independent Early Flowering) gene families are mediating to regulate FLC-related transcription factors by chromatic remodelling, are downregulated in controlgroup and upregulated in ricebean. TFL gene families reported from rice bean control inflorescence identity and maintain indeterminate inflorescences could be a better target for breeding purpose. These genes prevent the expression of APETALA1 and LEAFY, which play major roles in regulating the time of flowering. Regulations of FCA – Flowering time control, photoperiod-independent early flowering isoforms to reactivate FLC in early embryos and chromatin modelling on the basis of transcriptional regulation are the significant source for late flowering targets.

Palatability and digestibility-specific gene pooling increase the understanding of trait qualities that are preferred, including hard seededness, for all genotypes used by farmers in India. Taste modalities are aversive to each species and respondents. 37 genes are screened for palatability indexed ligands that are produced from 8 enzymatic responses. Expression of genes that are indexed for ligands bitterness are mapped with glycine max as control reference. We found expression of 15 genes in Gmax are down-regulated than ricebean are the scientific evident that these are genes the source of breeders to target and edit for palatable crops for domestic cultivars. Bitterness are elicited by diverge metabolite products from an enzymatic reaction. These metabolites are reported as bitter predictors and limit the threshold of receptor promiscuity. Out of reported 270 molecules as bitterness predictors, mung bean has 15 bitterness producible ligands that activates taste receptors – T2R in humans. These ligands are activated from 8 enzymatic reaction that are projected as chemosensory studied proteins for taste receptivity. Expressions of these 15 genes are indicative markers in Glycine max for bitterness. Anthranilate, chalcone, cyanidin, flavone, morin, palmitoyl, resveratrol and spermidine are the major functional molecule to determine palatability index of mung bean. These ligands are aversive to act as stimulus and opposed to olfactory. It produced from deamination of coregulated aromatic acids derived by subsequent hydrolytic and reduction by nonenzymatic compositions and storage conditions. Anthranilate N-methyltransferase-like, -synthase alpha subunit, -beta subunit and - phosphoribosyltransferase are upregulated in Glycine max that acts as taste modifiers (specific to Hypersensitivity Induced Reaction). Anthranililate basically involves in the synthesis of acridine alkaloids that activates most of (17988223). All other 28 genes, reported for palatability index are downregulated in Gmax and upregulated in mung bean. Pharmacophore of these 14 ligands are mapped closer proximity of flavonoid group. These promises, the evolution of these ligands from mapped genes are self-evident that C-ring of flavonoids are the precursive and crucial for taste receptor activation.

Ricebean genes prevails resistance to biotic and abiotic stress which is an issue that is generally addressed in most plant species in the process of domestication and crop improvement. In *Vigna umbellata*, we identified 97% of the coding domains of selected gene families such as those encoding the Rho-type GTPase-activating proteins, Serine-threonine protein kinases (TAO1), presynaptic morphology proteins and enhanced disease resistance 1 isoforms, which limit the initiation of cell death and establish a hypersensitive response.

Fourteen medicinally important genomes were compared with the genes of the rice bean assembly. Genes that are abundant provide in-depth transcript profiles of individual genes. This alignment facilitated the identification of candidate genes pertinent to the pathways of interest as well as non-pathway targets whose expression is consistent with the synthesis of medicinally valuable compounds in rice bean. *Rauwolfia serpentine* contains an ajmalan alkaloid-rich metabolite showing a high gene alignment frequency with rice bean. This metabolite is an anti-arrhythmic in the human system, and a mixture of rapidly interconverting epimers was found.

Critically, the potential of genetic improvement of rice bean can contribute to the sustainability of food production, nutrition, and feasible farmer-accessible flowering conditions, and consumer-accessible palatable crop. The draft of the *Vigna umbellata* genome provides genetic information for its domestication and indicates that this species is suitable for cultivation in large areas and potentially presents high commercial value. Because rice bean is closely related and similar to adzuki bean, mung bean and cowpea species, that we generated and analysed a high-density genetic map in rice bean, which could help to dissect important economic traits and meet the centuries-long challenge of its cultivation. This offers an avenue for scientists to improve the traits via CRISPR/Cas-Breeding and farmers to cultivate this legume crop variety harbouring immense nutritional potential for extensive consumption, to meet the centuries-long challenge of its cultivation and consumption. This is an effort to bring this nutritionally crucial legume crop in to mainstream commercial platform to serve several purposes including trait improvement for consumption, thereby alleviating the issue of micronutrient malnutrition, prevalent worldwide.

### Accession list of selected species under leguminous crops genome comparison

*Arachis hypogaea* : PIVG00000000.1; PRJNA476953, *Cajanus cajan* : AGCT00000000.1; PRJNA376605, *Cicer arietinum* : ANPC00000000.1; PRJNA190909, *Glycine max* : ACUP00000000.3; PRJNA19861,*Lupinus angustifolius* : CM007380.1; PRJNA356456, Mucuna pruriens : QJKJ00000000.1; PRJNA414658, *Phaseolus coccineus* : QBDZ00000000.1; PRJNA448610, *Phaseolus vulgaris* : ANNZ00000000.1; PRJNA240798, *Pisum sativum* : PUCA000000000.1; PRJNA432052, *Vicia faba* : CSVX00000000.1; PRJEB8906, *Vigna angularis*: JZJH00000000.1; PRJNA328963, *Vigna radiata* : JJMO00000000.1; PRJNA301363, and *Vigna unguiculata* : NBOW00000000.1; PRJNA521068.

*Vigna umbellata* assembled and annotated reads are deposited at NCBI Genome database. ID: SPEB00000000; PRJNA432818

## Acknowledgement

We sincerely thank to the Director of International Centre for Genetic Engineering and Biotechnology, New Delhi for providing fund from core fund to complete ricebean draft genome sequence.

